# Amyloid beta 42 alters cardiac metabolism and impairs cardiac function in obesity

**DOI:** 10.1101/2022.10.02.510555

**Authors:** Liam G Hall, Juliane K. Czeczor, Timothy Connor, Javier Botella, Kirstie A. De Jong, Mark C. Renton, Amanda J. Genders, Kylie Venardos, Sheree D. Martin, Simon T. Bond, Kathryn Aston-Mourney, Kirsten F. Howlett, James A Campbell, Greg R. Collier, Ken R. Walder, Matthew McKenzie, Mark Ziemann, Sean L. McGee

## Abstract

There are epidemiological associations between obesity and type 2 diabetes, cardiovascular disease and Alzheimer’s disease. While some common aetiological mechanisms are known, the role of amyloid beta 42 (Aβ_42_) in these diverse chronic diseases is obscure. Here we show that adipose tissue releases Aβ_42_, which is increased from adipose tissue of obese mice and is associated with higher plasma Aβ_42_. Increasing circulating Aβ_42_ levels in non-obese mice had no effect on systemic glucose homeostasis but had obesity-like effects on the heart, including reduced cardiac glucose clearance and impaired cardiac function. These effects on cardiac function were not observed when circulating levels of the closely related Aβ_40_ isoform were increased. Administration of an Aβ neutralising antibody prevented obesity-induced cardiac dysfunction and hypertrophy. Furthermore, Aβ neutralising antibody administration in established obesity prevented further deterioration of cardiac function. Multi-contrast transcriptomic analyses revealed that Aβ_42_ impacted pathways of mitochondrial metabolism and exposure of cardiomyocytes to Aβ_42_ inhibited mitochondrial function. These data reveal a role for systemic Aβ_42_ in the development of cardiac disease in obesity and suggest that therapeutics designed for Alzheimer’s disease could be effective in combating obesity-induced heart failure.

## MAIN

Patients with Alzheimer’s disease (AD) are predisposed to the development of cardiovascular disease^1^ and type 2 diabetes (T2D)^2^. Shared risk factors in the aetiology of these diseases include obesity, dyslipidemia, impaired glucose homeostasis and low aerobic fitness^3^. However, the pathogenic mechanisms involved remain unresolved. A potential molecular link is the amyloid beta (Aβ) peptide, which is a putative pathogenic driver of AD^4^.

Aβ peptides, ranging from 36-44 amino acids in length, are derived by proteolytic processing of the trans-membrane amyloid precursor protein (APP)^5^. Initial cleavage by either α- or β-secretases commits APP to the non-amyloidogenic or amyloidogenic pathways, respectively^5^. Following cleavage by the β-secretase BACE1, the remaining β-C-terminal fragment of APP can be cleaved by γ-secretase, which produces Aβ peptides that are released into the extracellular space via exocytosis^5^. The Aβ peptide of 42 amino acids (Aβ_42_) has a particular propensity to aggregate, form oligomers and higher order fibril structures that influence its physiological and pathological functions^4^. For example, Aβ_42_ can interact with and activate or inhibit numerous cell surface receptors, some of which include the N-methyl-D-aspartate (NMDA) receptor^6^, α-amino-3-hydroxy-5-methyl-4-isoxazolepropionic acid (AMPA) receptor^7^, metabotropic glutamate receptor 5 (mGluR5)^8^, β_2_-adrenergic receptor^9^, as well as pattern-recognition receptors such as Toll-like receptors^10^ and the receptor for advanced glycation end products (RAGE)^11^. Many of these receptor interactions are dependent on the oligomerisation state of Aβ_42_, which is highly dynamic and stochastic^11-13^. Extracellular Aβ_42_ can also be internalised where it can then interact with organelles, intracellular signalling molecules and enzymes, disrupting normal cellular function^14^. The dynamic and multidimensional structure of Aβ peptides, particularly Aβ_42_, means that dissecting their physiological and pathological functions has been challenging.

Accumulation of Aβ_42_ in the central nervous system is linked with alterations in metabolism, including impairments in glucose uptake^15,16^, glucose utilisation^17^ and aspects of mitochondrial function^18-21^ in several different cell types. Aβ_42_ also impairs neuronal insulin action by interacting with the insulin receptor^22^ and for this reason AD has been referred to as type 3 diabetes^4^. APP and its proteolytic processing pathways are expressed in peripheral tissues and Aβ peptides are found in plasma^23^. The expression of APP is increased in adipose tissue of obese humans^24^ and APP overexpression in adipose tissue of mice causes adiposity and insulin resistance due to impaired adipocyte mitochondrial function^25^. However, the role of circulating Aβ peptides in the regulation of peripheral metabolism remains largely unexplored. A recent study suggests that Aβ peptides released from peripheral tissues can inhibit insulin secretion from pancreatic beta cells^26^. Furthermore, plasma concentrations of Aβ_42_, but not Aβ_40_, positively correlate with fat mass in humans^27^, suggesting that systemic Aβ_42_ could play a role in the dysregulation of metabolism in obesity. The tissues that contribute to the increase in plasma Aβ_42_ in obesity and the metabolic consequences of elevated circulating Aβ_42_ remain unresolved and were examined in the present study.

## RESULTS

### Release of Aβ_42_ from adipose tissue is increased in obesity

To better understand the proteolytic processing of APP in the periphery in obesity, the expression and activity of APP and components of the amyloidogenic pathway were assessed in tissues from control and diet-induced obese mice. The expression of *App* was increased in adipose tissue of obese mice, but not in the liver or skeletal muscle (Fig. 1a). Similarly, *Bace1* (Fig. 1b) and *Psen1* (Fig. 1c), which encodes the presenilin 1 subunit of γ-secretase, were also increased in adipose tissue of obese mice, but not in the liver or skeletal muscle. The activity of BACE1 was increased in adipose tissue and reduced in the liver of obese mice (Fig. 1d). Adipose tissue explants were found to release Aβ_42_ but this was not different between control and obese mice when expressed relative to explant mass (Fig. 1e). However, when accounting for total fat pad mass, adipose tissue of obese mice released more Aβ_42_ (Fig. 1f). Release of Aβ_42_ was inhibited by incubation of explants with Brefeldin A (BFA; Extended Data Fig. 1a), an inhibitor of exocytosis^28^, indicating that adipose tissue actively secretes Aβ_42_. Consistent with greater Aβ_42_ release from adipose tissue of obese mice, the concentration of plasma Aβ_42_ was higher in obese mice (Fig. 1g). Similar to findings in humans^27^, plasma Aβ_42_ was significantly correlated with fat mass (Extended Data Fig. 1b) but not with lean mass (Extended Data Fig. 1c). Together these data show that Aβ_42_ is released from adipose tissue, which is increased in obesity.

**Figure 1:**
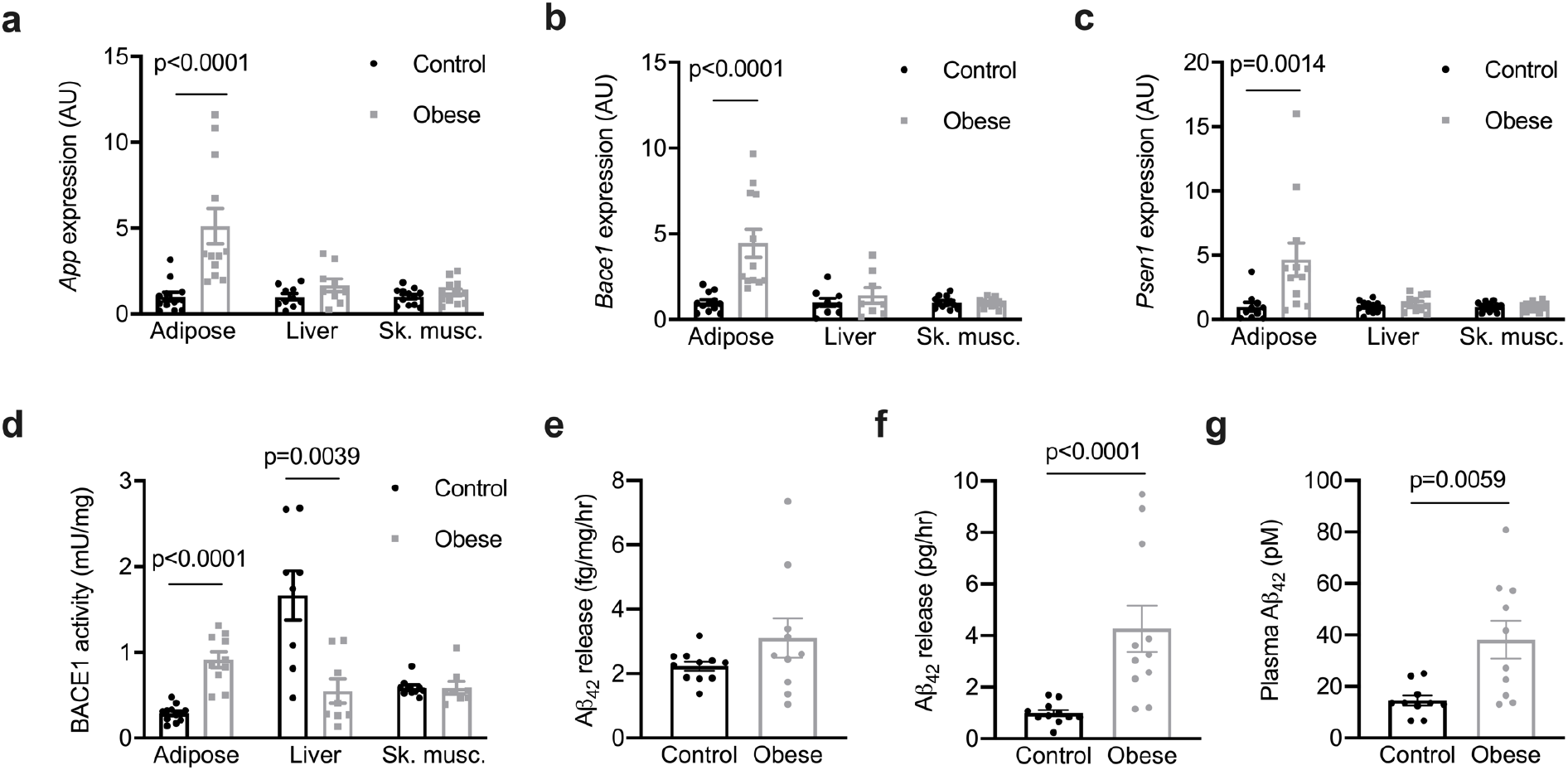
Aβ_42_ released from adipose tissue is increased in obesity. **a**, *App* expression in adipose tissue (Mann Whitney test, U=6), the liver and skeletal muscle (Sk. musc., quadriceps; unpaired t-tests) from control and obese mice. **b**, *Bace1* expression in adipose tissue (Mann Whitney test, U=1), the liver and sk. musc. (unpaired t-tests) from control and obese mice. **c**, *Psen1* expression in adipose tissue (Mann Whitney test, U=14) and, the liver and sk. musc. from control and obese mice (unpaired t-tests). **d**, BACE1 activity in adipose tissue, the liver and sk. musc. from control and obese mice (unpaired t-tests). **e**, relative release of Aβ_42_ from adipose tissue of control and obese mice normalised for tissue weight. **f**, absolute release of Aβ_42_ from adipose tissue of control and obese mice (Mann Whitney test, U=6). **g**, plasma Aβ_42_ in control and obese mice (unpaired t-test). Data are mean ± SEM, n = 10-12 mice per group. Statistical tests are two-tailed.

### Increasing systemic Aβ_42_ alters cardiac metabolism and impairs cardiac function

To examine the effect of increased circulating Aβ_42_ on systemic metabolism, mice were administered Aβ_42_ or a peptide corresponding to a scrambled Aβ_42_ sequence (ScrAβ_42_; 1μg/day i.p.) for 4 weeks (Fig. 2a). This administration regimen increased plasma Aβ_42_ ∼4-fold (Fig. 2b) but had no effect on body weight (Fig. 2c) or body composition (Extended Data Fig. 2a and b). Similarly, Aβ_42_ administration had no effect on blood glucose during both insulin (Fig. 2d) and glucose tolerance tests (GTT; Fig. 2e), or glucose-stimulated insulin secretion (Extended Data Fig. 2c). However, further analysis of glucose fate throughout the GTT using both 2-^2^H-deoxyglucose and 1-^14^C-glucose tracers revealed that glucose clearance by the heart was reduced in mice administered Aβ_42_ (Fig. 2f). In contrast, there were no differences in glucose clearance by skeletal muscle or adipose tissue (Extended Data Fig. 2d). The reduction in cardiac glucose clearance in mice administered Aβ_42_ was associated with additional alterations in cardiac glucose metabolism, including increased glucose incorporation into lipids (Fig. 2g), which was associated with a trend (p=0.0830) towards increased cardiac triglycerides (TG; Extended Data Fig. 2e). These alterations in cardiac metabolism in mice administered Aβ_42_ were independent of changes in plasma free fatty acids and lipids (Extended Data Fig. 2f-i). These data show that increasing systemic Aβ_42_ alters cardiac glucose metabolism.

**Figure 2:**
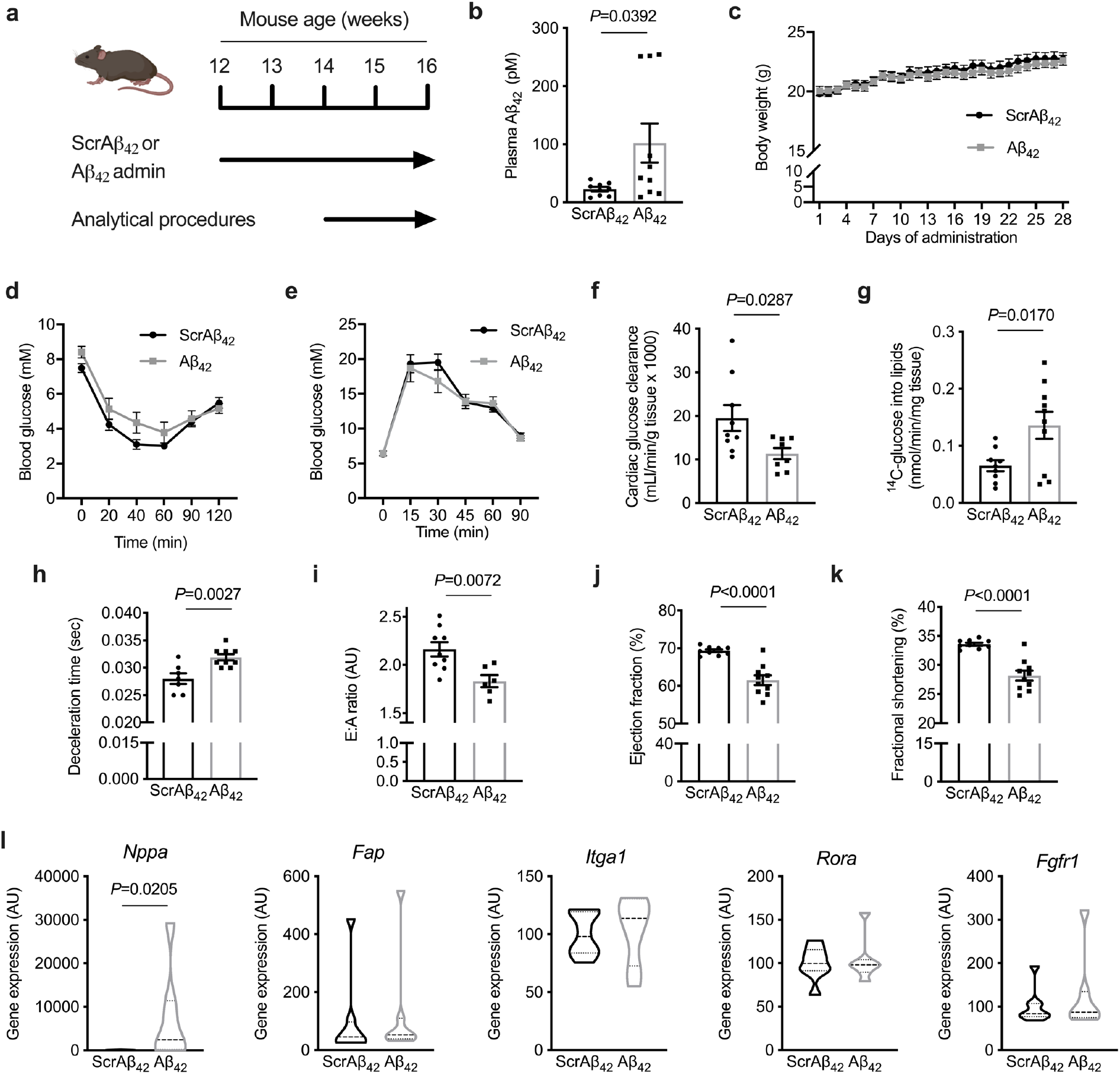
Aβ_42_ administration alters cardiac metabolism and impairs cardiac function. **a**, schematic of experiment where mice were administered Aβ_42_ or scrambled Aβ_42_ (ScrAβ_42_; 1μg/day i.p.) for four weeks and analytical procedures were performed in final two weeks. **b**, plasma Aβ_42_ in mice 5 hr after administration of ScrAβ_42_ or Aβ_42_ (unpaired t-test). **c**, body weight in mice administered ScrAβ_42_ or Aβ_42_. **d**, blood glucose during an insulin tolerance test in mice administered ScrAβ_42_ or Aβ_42_. **e**, blood glucose during a glucose tolerance test in mice administered ScrAβ_42_ or Aβ_42_. **f**, cardiac glucose clearance in mice administered ScrAβ_42_ or Aβ_42_ (unpaired t-test). **g**, ^14^C-glucose incorporation into lipids in mice administered ScrAβ_42_ or Aβ_42_ (unpaired t-test). **h**, deceleration time in mice administered ScrAβ_42_ or Aβ_42_ (unpaired t-test). **i**, E:A ratio in mice administered ScrAβ_42_ or Aβ_42_ (unpaired t-test). **j**, ejection fraction in mice administered ScrAβ_42_ or Aβ_42_ (unpaired t-test). **k**, fractional shortening in mice administered ScrAβ_42_ or Aβ_42_ (unpaired t-test). **l**, Expression of *Nppa* (Mann-Whitney test, U=8), *Fap, Itga1, Rora* and *Fgfr1* in hearts of mice administered ScrAβ_42_ or Aβ_42_. Data are mean ± SEM, n = 8-12 mice per group. Statistical tests are two-tailed.

Obesity results in reprogramming of cardiac metabolism that includes reduced glucose uptake and impaired glucose utilisation for a given insulin concentration^29^. These obesity-induced alterations in cardiac metabolism and resulting energetics have been linked to impairments in diastolic function that negatively impact cardiac relaxation and left ventricular (LV) filling^30^. Although increased filling pressures can overcome diastolic dysfunction in the early phases of the disease^31^, longer-term consequences include LV hypertrophy and eventual heart failure^31^. This form of heart failure most commonly manifests as heart failure with preserved ejection fraction (HFpEF)^31,32^. Hence, obesity-induced alteration of cardiac metabolism is one of the precipitating events that ultimately leads to obesity-induced heart failure^30,31^. As Aβ_42_ induced changes in cardiac metabolism that phenocopied those observed in obesity, the effect of Aβ_42_ on cardiac function was examined by echocardiography (Extended Data Fig. 2j). Administration of Aβ_42_ had wide-ranging effects on cardiac function, including impaired diastolic function, represented by increased deceleration time (Fig. 2h) and reduced E:A ratio (Fig. 2i), indicating impairments in cardiac relaxation. Furthermore, mice administered Aβ_42_ displayed evidence of impaired systolic function, including reduced ejection fraction (Fig. 2j) and fractional shortening (Fig. 2k). The estimated LV mass, calculated from M-mode measures, was lower in mice administered Aβ_42_ (Extended Data Table 1). To better understand how Aβ_42_ impairs cardiac function, profiling of genes that are differentially expressed in human heart failure in the major cell types of the heart^33^ was performed. In the hearts of mice administered Aβ_42_, there was a significant increase in the expression of *Nppa* (Fig. 2l), which encodes natriuretic peptide A, and is increased in cardiomyocytes in human heart failure^33^. In contrast, there were no changes in the expression of *Fap, Itga1, Rora* and *Fgfr1*, which are increased in heart failure in cardiac fibroblasts, pericytes, smooth myocytes and endothelial cells, respectively^33^. Together, these data indicate that Aβ_42_ impairs cardiac function, likely through effects on cardiomyocytes.

### Increasing systemic Aβ_40_ has no effect on cardiac function

In the context of AD, Aβ_42_ is considered the most pathogenic of the Aβ peptides, possibly because of its greater propensity to aggregate^34^. However, the abundance of Aβ_42_ in plasma is lower than Aβ _40_^23,35^. To determine whether Aβ _40_ also has deleterious effects on the heart, mice were administered Aβ_40_ (1μg/day i.p.) for 4 weeks followed by echocardiographic assessment of cardiac function and morphology, as was performed for Aβ_42_. Administration of Aβ_40_ markedly increased the circulating concentration of Aβ_40_ (Extended Data Fig. 3a) but had no effect on body weight and composition (Extended Data Fig. 3b-d), and had no effect on any index of cardiac function or morphology (Extended Data Table 2 and Extended Data Fig. 3e). Together with our previous findings, these data suggest that circulating Aβ_42_, but not Aβ_40_, has deleterious effects on the heart.

### An Aβ neutralising antibody prevents diastolic dysfunction in developing obesity

Having established that Aβ_42_ release from adipose tissue is increased in obesity and that raising systemic concentrations of Aβ_42_ negatively impacts cardiac metabolism and function, we next sought to determine whether Aβ_42_ mediates the adverse effects of obesity on cardiac function. We and others have previously established that deletion of *App* or components of the APP proteolytic machinery in mice increases energy expenditure and confers resistance to obesity^36,37^. Hence, these genetic models cannot be used to assess the role of Aβ_42_ on cardiac function in obesity. Instead, the Aβ neutralising antibody 3D6 was employed to address this question. Binding of 3D6 to Aβ_42_ prevents Aβ_42_ interacting with receptors and also cellular internalisation of Aβ_42_^38^. The humanised version of 3D6, termed bapineuzumab, reached phase III trials for AD^39^. Although it had excellent target engagement and an acceptable safety profile, it failed to produce meaningful improvements in cognition in AD patients^39^. Cardiovascular conditions were exclusion criteria for this trial and many others for AD therapies, meaning that data on any potential cardiovascular effects of Aβ_42_ are not available. Therefore, to determine whether Aβ_42_ mediates the adverse effects of obesity on cardiac function, mice were fed a high fat diet and were simultaneously administered 3D6 or a control antibody (0.75mg/kg, i.p. once weekly) for a period of 4 months. Cardiac function and morphology were assessed by echocardiography at the beginning and conclusion of the experiment (Fig 3a). The initiating event in the development of obesity-induced cardiac dysfunction and eventual heart failure is impaired cardiac relaxation, which can be characterised by increased deceleration time^40-42^. We have previously observed impairments in deceleration time in this model between 3-4 months of high fat feeding^43^. Consistent with previous studies^44^, 3D6 administration did not reduce plasma Aβ_42_ concentration (Extended Data Fig. 4a) nor had any effect on body weight and composition (Extended Data Fig. 4b-d). Furthermore, 3D6 administration had no effect on insulin tolerance, glucose tolerance or plasma insulin during the GTT (Extended Data Fig. 4e-g). In mice administered control antibody, deceleration time increased over the course of the high fat feeding period (Fig. 3b). In contrast, deceleration time was not increased in mice administered 3D6 (Fig. 3b). The same result was observed when deceleration time was normalised as a percentage of the cardiac cycle (Extended Data Fig. 4h). Although initial estimated LV mass was not matched between groups, LV mass was increased throughout the study in mice administered control antibody, while LV mass was unchanged in mice administered 3D6 (Fig. 4c). This was also supported by a similar finding in left ventricular posterior wall thickness at diastole (LVPWd; Extended Data Table 3), an alternate index of LV mass^45^. There were no other differences in cardiac function and morphology observed between groups (Extended Data Table 3). These data suggest that systemic Aβ_42_ contributes to the development of obesity-induced cardiac dysfunction, characterised by impaired diastolic function and increased LV mass.

**Figure 3:**
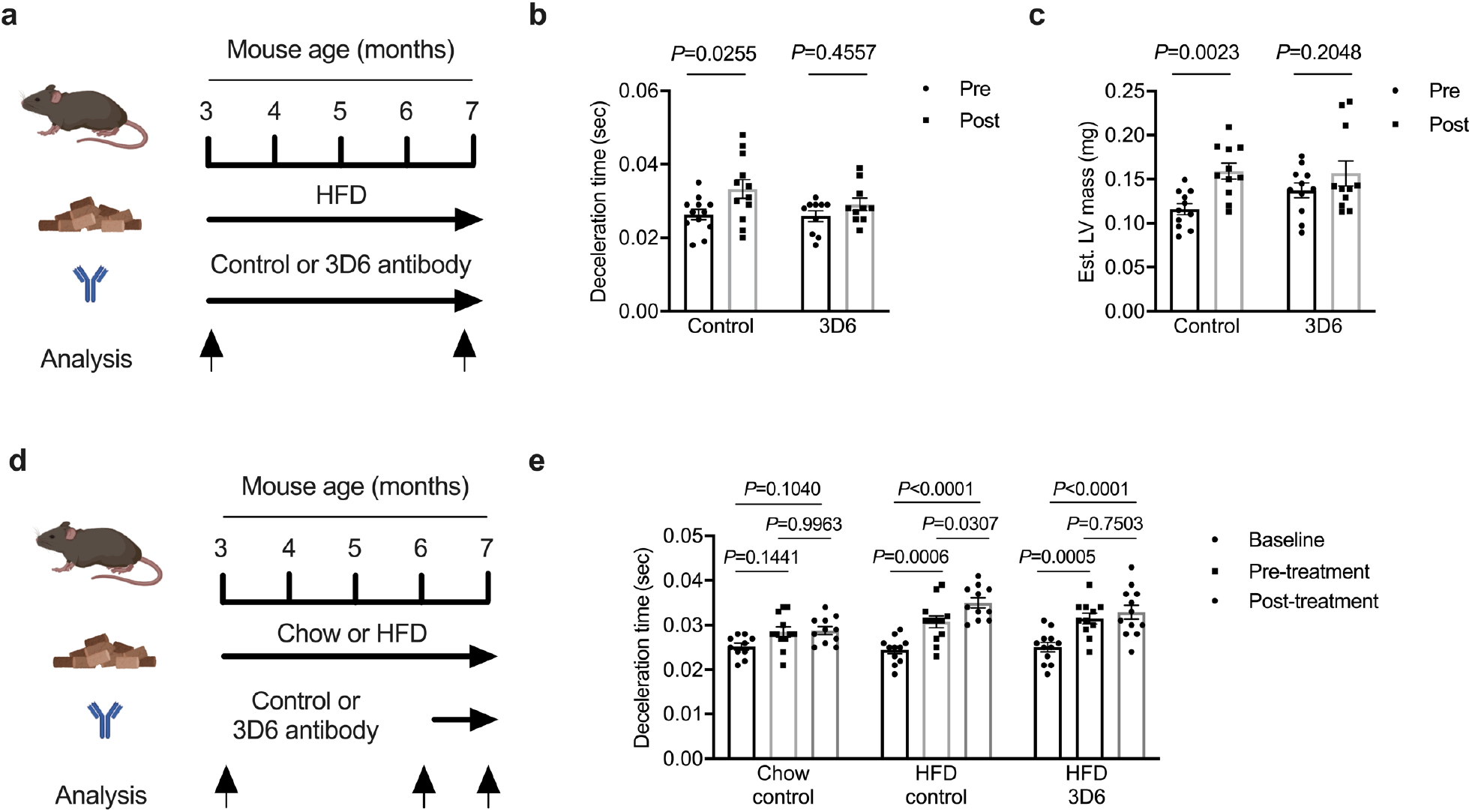
An Aβ_42_ neutralising antibody prevents diastolic dysfunction in developing obesity and prevents further diastolic decline in established obesity. **a**, schematic of experiment where mice fed a high fat diet (HFD) for 4 months and were simultaneously administered a control antibody or an Aβ_42_ neutralising antibody (3D6) once weekly. Analysis of cardiac function and morphology and function was performed at the beginning and end of the experiment. **b**, deceleration time in mice prior to and after 4 months of high fat feeding and antibody administration (two-way repeated measures ANOVA (time *P* = 0.0140, *F*(1,20)=7.257) and Sidak’s multiple comparisons test *P*.adjusted). **c**, estimated left ventricle (LV) mass in mice prior to and after 4 months of high fat feeding and antibody administration (two-way repeated measures ANOVA (time *P* = 0.0010, *F*(1,20)=14.93) and Sidak’s multiple comparisons test *P*.adjusted). **d**, schematic of experiment where mice fed either chow or a high fat diet (HFD) for 4 months and were administered a control antibody or an Aβ_42_ neutralising antibody (3D6) once weekly in the final month of the diet period. Analysis of cardiac function and morphology and function was performed at the beginning and of the experiment (baseline), prior to the treatment period (pre-treatment) and at the end of the treatment period (post-treatment). **e**, deceleration time in mice at Baseline, pre-treatment and post-treatment after 4 months of chow or high fat feeding and antibody administration (mixed-effects model (time *P* < 0.0001, *F*(2,93)=31.56; treatment *P* = 0.0151, *F*(2,93)=4.390) and Sidak’s multiple comparisons test *P*.adjusted). Data are mean ± SEM, n = 10-12 mice per group. Statistical tests are two-tailed.

**Figure 4:**
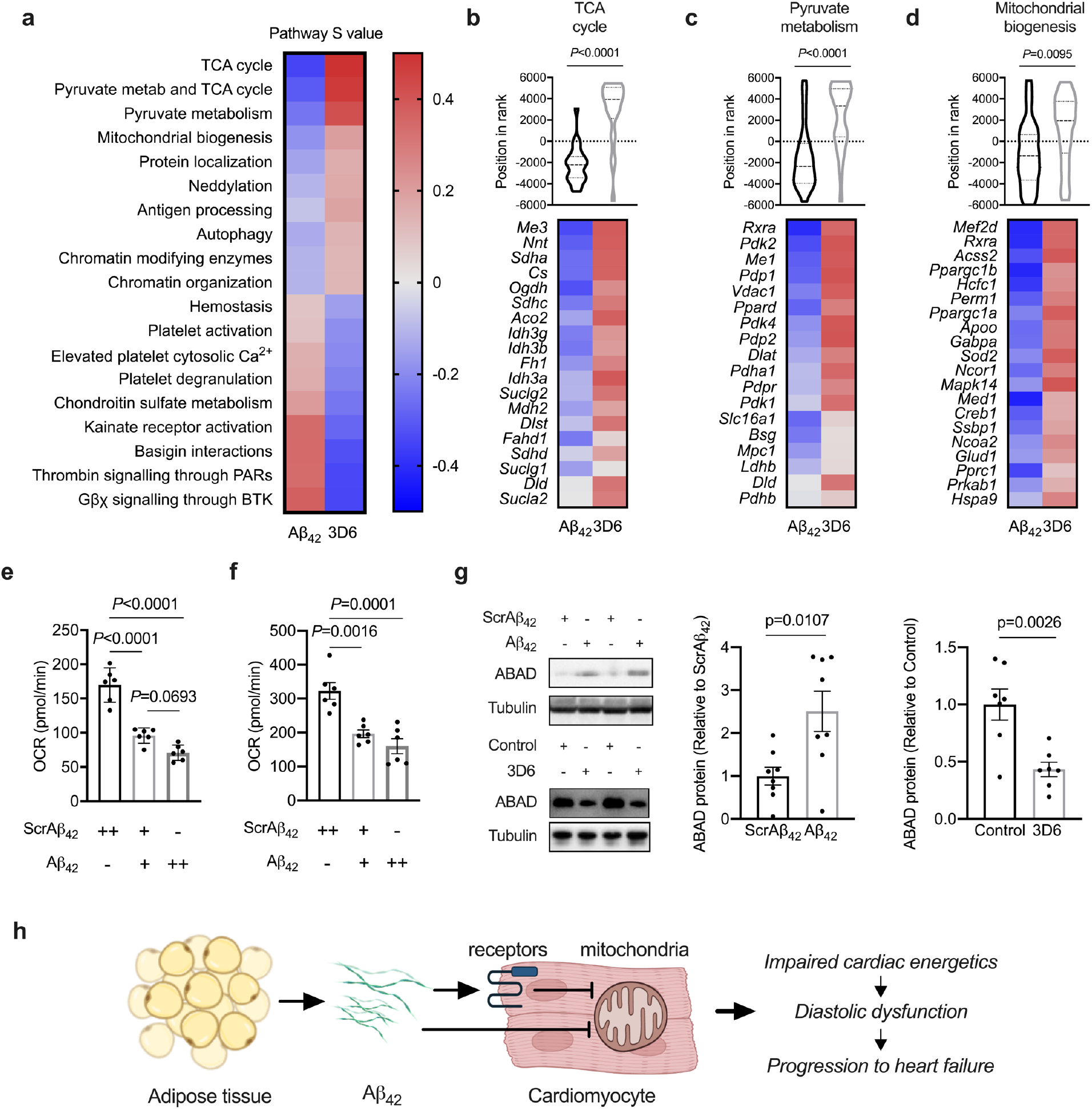
Aβ_42_ impairs cardiomyocyte mitochondrial function. **a**, heat map of Reactome pathways reciprocally regulated in the hearts of mice from Aβ_42_ administration (Fig. 2a) and 3D6 (Fig. 3a) administration studies, determined by MITCH analysis from bulk RNA-seq data. **b**, TCA cycle pathway rank (*P*.adjusted MANOVA test) and heat map of TCA cycle genes in the hearts of mice from Aβ_42_ administration and 3D6 administration studies. **c**, pyruvate metabolism pathway rank (*P*.adjusted MANOVA test) and heat map of pyruvate metabolism genes in the hearts of mice from Aβ_42_ administration and 3D6 administration studies. **d**, mitochondrial biogenesis pathway rank (*P*.adjusted MANOVA test) and heat map of mitochondrial biogenesis genes in the hearts of mice from Aβ_42_ administration and 3D6 administration studies. **e**, basal oxygen consumption (OCR) in primary mouse neonatal cardiomyocytes (NVCM) exposed to ScrAβ_42_ or Aβ_42_ for 48 hrs (one-way ANOVA (*P* < 0.0001; F(2,15) = 53.5) with Sidak’s repeated measures test *P*.adjusted). **f**, maximal OCR in primary NVCM exposed to ScrAβ_42_ or Aβ_42_ for 48 hrs one-way ANOVA (*P* = 0.0001; F(2,15) = 17.7) with Sidak’s repeated measures test *P*.adjusted). **g**, amyloid binding alcohol dehydrogenase (ABAD) abundance in the hearts of mice from Aβ_42_ administration and 3D6 administration studies. Data are mean ± SEM, n = 7-8 mice per group, n = 6 biological replicates per group for cardiomyocyte analyses. Statistical tests are two-tailed. **h**, schematic of proposed model whereby adipose tissue release of Aβ_42_ is increased in obesity resulting in higher circulating levels of Aβ_42_. Increased circulating Aβ_42_ inhibits cardiomyocyte mitochondrial function and causes diastolic dysfunction, which starts the progression towards heart failure. The effects of Aβ_42_ on cardiomyocytes could be mediated through receptor-mediated signalling, receptor-mediated internalisation, or direct internalisation.

### An Aβ neutralising antibody prevents further diastolic decline in established obesity

At present there are limited effective therapies to treat cardiac dysfunction in obesity that prevent the decline to heart failure. While most therapies are focussed on reducing hypertension, it is accepted that cardiac dysfunction in obesity can develop independently of hypertension^46^. Our finding that Aβ_42_ antagonism prevents obesity-induced impairment of cardiac relaxation raises the possibility that targeting Aβ_42_ could be an effective treatment for obesity-induced cardiac dysfunction. To test this hypothesis, mice were fed either standard chow or a high fat diet for 4 months and were administered 3D6 or control antibodies for the final 4 weeks of the experiment. Echocardiography to assess cardiac function and morphology was performed at the beginning of the experiment (baseline), prior to antibody treatment (pre-treatment) and at the conclusion of the study (post-treatment; Fig 3d). Mice fed a high fat diet had increased body weight, fat mass and fasting plasma insulin compared with chow fed mice, however 3D6 administration had no effect on these parameters (Extended Data Fig. 5a-e). There was no change in deceleration time in chow fed mice administered control antibody throughout the experiment (Fig. 3e). Mice fed high fat diet and administered control antibody had an increase in deceleration time from baseline to pre-treatment, which increased further from pre-treatment to post-treatment (Fig. 3e). Mice fed high fat diet and administered 3D6 antibody had an increase in deceleration time from baseline to pre-treatment, however there was no further increase in deceleration time from pre-treatment to post-treatment (Fig. 3e). These findings were consistent when deceleration time was expressed as a percentage of the cardiac cycle (Extended Data Fig. 5f). Short term administration of 3D6 in established obesity did not alter any index of LV mass (Extended Data Table 4). These data suggest that Aβ_42_ antagonism halts further deterioration of cardiac function in established obesity.

### Aβ_42_ induces mitochondrial dysfunction in cardiomyocytes

In the context of AD, Aβ_42_ has highly complex cellular effects through a myriad of mechanisms depending on the target cell type and its aggregation state^47^. One of the mechanisms by which Aβ_42_ impairs metabolism in AD is through competition with insulin for binding to the insulin receptor^22^. Although this occurs at Aβ concentrations orders of magnitude higher than circulating Aβ_42_ concentrations^22^, accumulation of Aβ_42_ at certain tissues can reach these levels^23^. To assess whether increased circulating Aβ_42_ impacts cardiac insulin signalling, the phosphorylation of Akt at its two regulatory sites was examined in heart samples collected immediately after the glucose tolerance test in Aβ_42_ administration studies (Fig. 2a). No differences in Akt phosphorylation between groups were observed (Extended Data Fig. 6a), suggesting that other mechanisms are involved.

To gain unbiased insights into the deleterious effects that Aβ_42_ has on the heart, a multidimensional transcriptomic approach was employed. Transcriptomic data from the LV of hearts obtained in Aβ_42_ administration studies (Fig. 2a) were compared with data obtained in the 3D6 administration study (Fig. 3a) and pathways reciprocally regulated in these experiments were identified using mitch^48^. It was reasoned that reciprocally regulated pathways from these experiments represent the genesets fundamental to the effects of Aβ_42_ on the heart. Global gene expression was evenly distributed across the four quadrants of the two-dimensional analysis, whether represented by input profile or by gene ranking (Extended Data Fig. 6b and c). Multi-contrast enrichment analysis identified the TCA cycle, pyruvate metabolism and mitochondrial biogenesis pathways with the largest effect sizes that were reduced by Aβ_42_ administration and increased by 3D6 administration (Fig. 4a and Extended Data Table 5). These pathways are dysregulated in obesity-induced cardiac dysfunction and have been linked with disrupted energetics that cause impaired cardiac relaxation^30,49-51^. The transcriptional reprogramming of the TCA cycle included altered regulation of genes encoding TCA cycle enzymes (Fig. 4b), while transcriptional regulation of pyruvate metabolism included subunits of pyruvate dehydrogenase (PDH) and PDH kinases (PDKs; Fig 4c). Regulated genes in the mitochondrial biogenesis pathway were mainly of nuclear transcription factors controlling the mitochondrial biogenesis process (Fig. 4d). Interestingly, several receptor pathways were increased by Aβ_42_ administration and reduced by 3D6 administration (Fig. 4a and Extended Data Table 5).

As our previous expression profiling suggested that the effects of Aβ_42_ were specific for cardiomyocytes, the effect of Aβ_42_ on cardiomyocyte mitochondrial respiration was examined. Primary neonatal ventricular cardiomyocytes (NVCM) were exposed to increasing concentrations of Aβ_42_ for 48hrs, which reduced both basal respiration (Fig. 4e) and maximal respiratory capacity (Fig. 4f). This was not explained by a loss of cell viability (Extended Data Fig. 6d) and indicate that Aβ_42_ has direct effects on cardiomyocytes. The widespread transcriptional reprogramming of mitochondrial pathways likely indicates that this is a secondary response to effects of Aβ_42_ on oxidative metabolism. A well-characterised mechanism by which Aβ peptides induce mitochondrial dysfunction is through cellular internalisation and import into mitochondria^52^. A mitochondrial target of Aβ peptides is the amyloid beta binding alcohol dehydrogenase (ABAD), a short-chain dehydrogenase that oxidises a broad change of substrates and is an important enzyme in the β-oxidation of fatty acids^53^. Binding of Aβ causes a change in ABAD confirmation that reduces its enzymatic activity towards certain substrates, increases ABAD abundance and results in mitochondrial dysfunction^52^. Cardiac ABAD abundance was increased in mice administered Aβ_42_ and was reduced in obese mice administered 3D6 (Fig. 4g). In Aβ_42_ administration studies, ABAD was not increased in any other tissue (Extended Data Fig. 6e). These findings suggest that systemic Aβ_42_ is internalised by cardiomyocytes and taken up into mitochondria, where it can influence mitochondrial function as well as impact on mitochondrial transcriptional programs.

## DISCUSSION

Our findings have revealed an unexpected role for Aβ_42_ in the aetiology of obesity-induced cardiac dysfunction. These data point to a paradigm where increased release of Aβ_42_ by adipose tissue and increased circulating Aβ_42_ in obesity negatively impacts cardiomyocyte mitochondrial function and causes cardiac dysfunction, particularly impaired cardiac relaxation, which is the precipitating event leading to heart failure (Fig. 4h). The association between increasing adiposity and risk of heart failure is well established, however the exact mechanisms involved remain poorly defined. This study identified Aβ_42_ as a factor linking adipose tissue expansion with impaired cardiac function. Furthermore, our data indicate that Aβ_42_ is a tractable therapeutic target for obesity-induced cardiac dysfunction.

Circulating concentrations of Aβ_42_ are correlated with fat mass^27^ and App expression is increased in adipose tissue of obese humans^24^. While these findings could indicate increased production and release of Aβ_42_ by adipose tissue, another plausible explanation is that Aβ_42_ clearance is reduced in obesity. Both Aβ_42_ and insulin are degraded by insulin degrading enzyme (IDE)^54^ and as both fasted and post-prandial insulin levels are generally higher in obesity, this could result in competition for IDE activity and reduced Aβ_42_ degradation. Indeed, plasma insulin and Aβ_42_ are highly correlated in humans^27^. However, our data show that the expression and activity of the amyloidogenic proteolytic machinery are increased in adipose tissue in obesity. Furthermore, release of Aβ_42_ by adipose tissue is increased in obesity and is directly related to adipose tissue mass. The expression and activity of BACE-1, which is obligatory for Aβ generation, appears to be a critical node controlling adipose Aβ_42_ release. Inflammatory signalling increases Bace-1 expression and activity in neurons, while post-translational modifications (PTMs) acutely regulate BACE-1 activity^55^. These include metabolically sensitive PTMs such as glycosylation and palmitoylation^55^. This suggests that the regulation of BACE-1 expression and activity, and Aβ_42_ release by adipose tissue may be intrinsically linked to metabolic perturbations and inflammation, which are hallmark features of obesity.

Our findings are consistent with associative observations linking Aβ_42_ to alterations in cardiac and cardiomyocyte function. For example, in patients with AD, Aβ_42_ accumulates in the heart and is associated with diastolic dysfunction^56^. Elevated circulating Aβ_42_ and diastolic dysfunction are also associated in persons with Down syndrome^57,58^. Furthermore, exposure of isolated rat hearts to Aβ_42_, albeit at supraphysiological levels, results in disrupted cardiac contractility^59^. Our multi-contrast transcriptomics analysis suggests that dysregulated mitochondrial function is central for the effects of Aβ_42_ on the heart and that Aβ_42_ targeting of mitochondrial ABAD could be an important contributor to obesity-induced mitochondrial dysfunction. Binding of Aβ_42_ to ABAD, also known as 3-hydroxyacyl-CoA dehydrogenase (HADH2), results in a rapid increase in ABAD abundance^52^, which has been observed in unbiased proteomic examinations of the heart in obesity^60,61^. In these studies, an increase in ABAD was associated with reduced respiratory responses in both sub-sarcolemmal and inter-myofibrillar mitochondria^60^ and increased injury in response to ischemia/reperfusion^61^. Furthermore, exposure of isolated rat heart mitochondria or immortalised H9C2 cardiomyoblasts to Aβ_42_ causes mitochondrial dysfunction, not only affecting energetics but also resulting in elevated ROS production, a loss of mitochondrial membrane potential, mitochondrial swelling, and impaired capacity to retain calcium^62^. In contrast, Aβ_40_ had less pronounced effects on these parameters of mitochondrial function^62^. Together, these findings support our conclusion that the effects of Aβ_42_ on the heart are in part due to the internalisation of Aβ_42_, but not Aβ_40_, by cardiomyocytes. The mechanisms of Aβ_42_ internalisation by cardiomyocytes remain unclear. In neuronal cells, oligomeric Aβ peptide internalisation can be mediated by macropinocytosis that requires both lipid rafts and active heparan sulfate proteoglycans^63^. Interestingly, the closely related chondroitin sulfate metabolism pathway was reciprocally regulated in our multi-contrast transcriptomic analysis (Fig. 4a and Extended Data Table 5). Furthermore, Aβ uptake can also occur through receptor-mediated internalisation and a number of different receptors have been implicated^64^, some of which were also identified in our transcriptomics analysis (Fig. 4a and Extended Data Table 5). Whether these receptor classes are also actionable targets for obesity-induced cardiac dysfunction remains to be determined. However, given the propensity of aggregated Aβ_42_ to associate with multiple receptors, it is likely that any receptor antagonist-based approach might have to include multiple therapies.

Findings from the present study add to a growing body of evidence indicating physiological and pathological roles for Aβ peptides in the periphery. The observation that Aβ_42_ administration induced both diastolic and systolic dysfunction, yet only diastolic dysfunction is observed in diet-induced obese mice, is likely explained by the complex physiology of obesity that include metabolic alterations mediated by hyperinsulinemia, other hormonal factors and increased cardiac substrate availability, which together could preserve systolic function. Nonetheless, our findings support the idea that therapies originally designed for AD that reduce or antagonise Aβ_42_ could be repurposed to treat and/or prevent obesity-induced heart failure, particularly those with HFpEF, for which treatment options are limited^31^. This is an urgent unmet need as this patient group has a 5-year mortality rate of up to 75% and currently affects up to 8% of people in certain populations^65^. Furthermore, the prevalence of HFpEF is predicted to rise in parallel with rising rates of obesity^65^. A suite of biologics and small molecules have been developed for AD that antagonise or reduce Aβ_42_, many with well-established safety and tolerability profiles in humans. These diverse therapies with distinct mechanisms of action could therefore be rapidly assessed as new treatments for heart failure in obesity.

In conclusion, this study has identified that Aβ_42_ release from adipose tissue is increased in obesity and is associated with elevated circulating Aβ_42_ and impairments in cardiac metabolism and function. Antagonism of Aβ_42_ prevented the development of obesity-induced cardiac dysfunction and slowed the progression of cardiac dysfunction in established obesity. Therefore, these findings support further studies assessing the repurposing of AD therapies for obesity-induced heart failure, which could fill a devastating therapeutic gap.

## METHODS

### Animal studies

Male C57BL6 mice were obtained from the Animal Resources Centre (Perth, WA, Australia) and were group housed with a 12 hr light/dark cycle at 22°C. Animals had ad libitum access to food and water. All procedures involving animals were approved by The Deakin University Animal Welfare Committee (A58-2010, G07-2013, G15-2017 and G08-2020), which is subject to the Australian Code for the Responsible Conduct of Research.

#### High fat feeding

At 12 weeks of age, mice were randomly assigned to regular chow or high fat diet (HFD; 43% digestible energy from fat, 20% from sucrose; SF04-001 Specialty Feeds). Throughout the diet period, mice were weighed weekly. After three months of the diet period, mice underwent body composition assessment using EchoMRI. Two days later, mice were fasted for 5 hrs before being humanely killed by cervical dislocation. Epididymal and inguinal adipose tissues were carefully excised and weighed and were either immediately frozen and stored at -80°C for later analysis or were used for *ex vivo* adipose explant experiments.

#### Aβ peptide administration

Lyophilised recombinant Aβ_42_ (#AG912, Millipore) and scrambled control peptide (ScrAβ_42_; #AG916, Millipore) were resuspended in 1% NH_4_OH and aliquoted at 200ng/ml in H_2_O and stored at -80^°^C for no longer than 4 weeks. Recombinant human Aβ_42_ was used as it has a greater propensity to aggregate than the mouse sequence^66^ and peptides appeared as low molecular weight aggregates of Aβ_42_ by SDS-PAGE (data not shown). Peptide administration began when male C57BL6J mice were 12 weeks of age and involved a daily i.p. injection of 1μg of peptide. Throughout the peptide administration period, body weight was measured daily and food intake was measured twice weekly, and body composition was measured weekly by EchoMRI. After two weeks, plasma Aβ_42_ was determined from samples collected 5 hr after recombinant ScrAβ_42_ or Aβ_42_ administration. After 3 weeks, an insulin tolerance test (ITT; i.p. 0.75U/kg) was performed following a 5 hr fast. Blood glucose was measured at baseline and after 20, 40, 60, 90 and 120 min after glucose administration. One week later, a glucose tolerance test (GTT; i.p 2g/kg with 5μCi ^3^H-2-deoxyglucose and 5μCi ^14^C-glucose) was performed after a 5 hr fast. Blood glucose was measured at baseline and after 15, 30, 45, 60 and 90 min after glucose administration. Plasma was obtained at 0, 15, 30, 60 and 90 min for insulin determination and the measurement of tracer specific activity. Mice were humanely killed by cervical dislocation immediately after the GTT. Blood was collected for plasma and tissues were dissected, rapidly frozen and stored at -80°C for later analysis. Another cohort of mice were administered ScrAβ_42_ or Aβ_42_ for four weeks, as described above. After four weeks of peptide administration, mice were anaesthetised by isoflurane inhalation before undergoing echocardiography. Two days later, mice were humanely killed by cervical dislocation and blood was obtained for plasma and tissues were dissected, rapidly frozen and stored at -80°C for later analysis. Another cohort of mice were administered ScrAβ_40_ or Aβ_40_ (Millipore) for four weeks, as described above. After four weeks of peptide administration, mice were anaesthetised by isoflurane inhalation before undergoing echocardiography. Two days later, mice were humanely killed by cervical dislocation and blood was obtained for plasma and tissues were dissected, rapidly frozen and stored at -80°C for later analysis.

#### 3D6 administration

For the 3D6 prevention study, 12-week-old mice were assessed for baseline cardiac structure and function by echocardiography and were assigned to one of two groups such that deceleration time was matched. One week later, mice were fed HFD (as described above) and then received control antibody (IgG2a isotype antibody, #BE-0085 BioXCell) or 3D6 antibody (#TAB-0809CLV, Creative Biolabs) weekly via i.p. injection (0.75mg/kg). Body weight was measured twice per week, while body composition was measured every two weeks throughout the study. After 10 weeks of high fat feeding, mice underwent an ITT and then a GTT after 11 weeks of high fat feeding. After 12 weeks of high fat feeding mice underwent echocardiography and two weeks later were humanely killed by cervical dislocation following a 5 hr fast. Blood was collected for plasma and tissues were dissected, rapidly frozen and stored at -80°C for later analysis. In the 3D6 reversal study, 12-week-old mice were assessed for baseline cardiac structure and function by echocardiography and were assigned to one of three groups such that deceleration time was matched. One week later, one group of mice remained on standard chow, while the other two groups were fed a high fat diet. Body weight was measured twice per week throughout the study. After 13 weeks of high fat feeding, mice underwent echocardiography for pre-treatment assessment of cardiac function and morphology. Mice were then administered control antibody or 3D6 antibody weekly via i.p. injection (0.75mg/kg) for the remainder of the study. Four weeks later, mice underwent post-treatment assessment of cardiac function and morphology, followed by assessment of body composition by EchoMRI two days later. One week later, mice were humanely killed by cervical dislocation following a 5 hr fast. Blood was collected for plasma and tissues were dissected, rapidly frozen and stored at -80°C for later analysis.

#### Echocardiography

Echocardiography using a Phillips HD15 diagnostic ultrasound system with 15 MHz linear-array transducer was performed as previously described^43,67,68^. In line with published recommendations^69^, depth of anaesthesia was modulated to maintain heart rate between 400 and 650 bpm. Transmitral Doppler imaging was used to assess LV filling velocity. Aortic valve Doppler imaging was used to assess aortic flow and ejection times. Ejection fraction and fractional shortening (FS) were derived from M-Mode measurements as indicators of LV systolic and contractile function and chamber morphology was also determined from M-mode measures, as previously described^68^. Echo images were analysed using the HD15 system in a blinded manner.

### Adipose tissue explants

Prior to the *ex vivo* incubation protocol, all consumables such as 12-well plates, tips, and tubes were blocked in 5% bovine serum albumin (BSA) to minimise binding of Aβ_42_ to plasticware^70^. Tests with recombinant Aβ_42_ revealed recovery of ∼95% of Aβ_42_ using this method. Upon collection of inguinal adipose tissue samples, four ∼10-15 mg tissue portions were isolated and each of them was placed in a well with 500 uL of DMEM media supplemented with 10% BSA. Three samples were incubated in vehicle (0.1% DMSO), while one incubated in 10μM Brefeldin A. Each tissue was individually weighed, the starting time of the incubation recorded, and the plate was placed at 37°C in 10% CO_2_ for 24 h. Following the 24 h period, both media and the tissue were rapidly collected, and subsequently snap-frozen, for further analyses. High sensitivity ELISA kits were used to quantify Aβ_42_ (#292-64501, FUJIFILM Wako) and the protocol was followed as per the manufacturer’s protocol. In brief, samples were brought to room temperature, vortexed, and 100 μl of sample was placed in each well, together with the respective internal controls for each plate. Each plate was then sealed and refrigerated overnight. Following this, the solutions were discarded, and each well was washed 5 times before addition of 100 μl of HRP-conjugated antibody. Following 1 hour of incubation, the solution was discarded and the washing step was repeated, with subsequent addition of 100 μl TMB solution to initiate the HRP reaction at room temperature in the dark. Following 30 minutes, 100 μl of Stop-solution was added to terminate the reaction and the absorbance (at 450nm) was subsequently measured to determine the protein levels.

### Glucose clearance and fate

The LV of the heart, quadriceps skeletal muscle and epididymal adipose tissue (15-25mg) were used to determine 2-^2^H-deoxyglucose clearance as we have previously described^68^. To determine 1-^14^C-glucose incorporation into glycogen, ∼10-15mg of LV tissue was digested in 1M KOH at 70°C for 20 min and glycogen was precipitated with saturated Na_2_SO_4_, washed twice with 95% ethanol and resuspended in acetate buffer (0.84% sodium acetate, 0.46% acetic acid, pH 4.75) containing 0.3mg/mL amyloglucosidase (Sigma). Glycogen was allowed to digest overnight at 37°C before being assayed for glucose content using the glucose oxidase method^71^. Digested glycogen was also assessed for ^14^C-glucose incorporation by scintillation counting. To determine 1-^14^C glucose incorporation into lipids, 5-10mg of LV tissue was homogenised in chloroform/methanol (2:1) and mixed overnight at room temperature. Organic and inorganic phases were separated by addition of 0.6% NaCl and the lower organic phase was collected and evaporated under N_2_ at 45°C. The dried extract was resuspended in absolute ethanol and TG content was assayed using TG GPO-PAP reagent (Roche) and ^14^C-glucose incorporation by scintillation counting.

### Gene expression and analysis

Bulk RNA was isolated from LV tissue by homogenisation using a hand-held homogeniser in Trizol and RNA was isolated using RNeasy kits (Qiagen), according to manufacturer’s instructions. Sequencing libraries were generated from 0.5 µg of total RNA using TruSeq Stranded Total RNA preparation kit (Illumina) as per manufacturer’s instructions. Transcriptome-wide mRNA levels were measured using the NovaSeq 6000 Sequencing System (Illumina). Data analysis was performed as previously described^72^. Briefly, sequence Fastq files underwent quality trimming and mapping to the mouse reference transcriptome with Salmon. The reference transcriptome sequence was obtained from Gencode (version vM24). Counts were read into R (version 4.0.2). Differential expression analysis was performed separately for two contrasts (ScrAβ_42_ vs. Aβ_42_ and Control vs 3D6). Genes with fewer than 10 reads per sample on average were discarded. Differential analysis was performed with DESeq2 (version1.28.1). Multi-contrast enrichment analysis was performed with mitch (version 1.0.6)^48^. The expression of *App, Bace1 and Psen1* were quantified in epididymal adipose tissue, quadriceps skeletal muscle and the liver of control and obese mice by real time RT-PCR using the following primer pairs (*App* Fwd CTTGCACGACTATGGCATGC, rev GTCATCCTCCTCTGCATCCG; *Bace1* Fwd GGAGCCCTTCTTTGACTCCC, rev CCCGTG TATAGCGAGTGGTC; *Psen1* Fwd TTCAAGAAAGCGTTGCCAGC, rev AGGGCTGCACA AGGTAATCC).

### Plasma and biochemical analyses

BACE-1 activity was assessed using a fluorescent assay kit (Merck), while plasma glycerol and plasma HDL-C were determined using colorimetric kits (Sigma), all according to manufacturer’s instructions. High sensitivity ELISAs were used to measure plasma Aβ_42_ and Aβ_40_ (FUJIFILM Wako), as described above, and plasma insulin (ALPCO), according to manufacturer’s instructions. Plasma TG concentration was assayed using TG GPO-PAP reagent (Roche) and plasma NEFA was determined using a colorimetric kit (FUJIFILM Wako).

### Primary cardiomyocytes

Neonatal ventricular cardiomyocytes were isolated from mixed sex, two-day-old C57BL6 mice as previously described^73^. Cells were seeded into Seahorse V7 assay plates at 60,000 cells/well. Prior to assays, cells were exposed to Aβ_42_ at 0, 200 and 300pM and co-incubated with ScrAβ_42_ at 300, 100 and 0pM, respectively, such that all cells were exposed to 300pM of total peptide for 2 days, with media replenished every 24 hr. These supraphysiological Aβ_42_ concentrations were used as we measured Aβ_42_ levels in FBS containing media at ∼20pM. Seahorse analyses were performed as previously described^74^. Cell viability was assessed by measuring extracellular and intracellular lactate dehydrogenase (LDH) following exposure of cardiomyocytes to 300pM of ScrAβ_42_ or Aβ_42_ for 48 hrs. LDH was measured as previously described^71^.

### Western blotting

Western blotting was performed as previously described^75^ using antibodies against pT308 Akt, pS473 Akt, total Akt (Cell Signalling Technology) and ABAD (MyBioScource).

### Statistical analyses

All data are presented as mean ± SEM and were assessed for normality using a Shapiro-Wilks test. Normally distributed data were analysed by independent samples t-test, one-way ANOVA, two-way ANOVA, two-way repeated measures ANOVA or mixed effects model, as described for statistically significant data in relevant figure legends. Data not normally distributed were analysed by Mann-Whitney test or Kruskal-Wallis test, as described for statistically significant data in relevant figure legends. All statistical analyses were performed using GraphPad Prism 8, with p values < 0.05 considered significant.

## ACKNOWLEDGEMENTS

The authors wish to thank Dr Tom Gumley, Assoc. Prof. John Amerena, Prof. Mark Hargreaves, Dr Sophie Hussey and Professor Roberto Cappai for their assistance with this study. The authors also wish to thank members of the Metabolic Research Unit for helpful discussions and Dr Richard Woolley for his assistance with echocardiography. The authors would also like to thank the Deakin University Animal Services team for their assistance throughout this study. Aspects of this work were funded by grants from the National Health and Medical Research Council (GNT1027226), the Diabetes Australia Research Program (Y17G-MCGS), the Ramaciotti Foundation and Ambetex Pty Ltd to SLM.

## AUTHOR CONTRIBUTIONS

Conceptualization, LGH, JKC and SLM; Methodology, LGH, JKC, TC, JB, KADJ, MCR, AJG, KV, SDM, STB, KAM, KFH, MM, MZ and SLM; Resources, KFH, JAC, GRC, KW, MM, MZ and SLM; Writing – original draft, SLM; Writing – review and editing, all authors; Funding acquisition, SLM.

## DATA AVAILABILITY

RNA-seq data generated in this study can be found at Gene Expression Omnibus, submission GSE213708.

## REFERENCES

1. Stampfer, M.J. Cardiovascular disease and Alzheimer’s disease: common links. Journal of internal medicine 260, 211–223 (2006).

2. Ott, A., et al. Diabetes mellitus and the risk of dementia: The Rotterdam Study. Neurology 53, 1937–1942 (1999).

3. Zhang, X.X., et al. The Epidemiology of Alzheimer’s Disease Modifiable Risk Factors and Prevention. J Prev Alzheimers Dis 8, 313–321 (2021).

4. Vignini, A., et al. Alzheimer’s disease and diabetes: new insights and unifying therapies. Current diabetes reviews 9, 218–227 (2013).

5. O’Brien, R.J. & Wong, P.C. Amyloid precursor protein processing and Alzheimer’s disease. Annu Rev Neurosci 34, 185–204 (2011).

6. Snyder, E.M., et al. Regulation of NMDA receptor trafficking by amyloid-beta. Nat Neurosci 8, 1051–1058 (2005).

7. Hsieh, H., et al. AMPAR removal underlies Abeta-induced synaptic depression and dendritic spine loss. Neuron 52, 831–843 (2006).

8. Um, J.W., et al. Metabotropic glutamate receptor 5 is a coreceptor for Alzheimer abeta oligomer bound to cellular prion protein. Neuron 79, 887–902 (2013).

9. Wang, D., Yuen, E.Y., Zhou, Y., Yan, Z. & Xiang, Y.K. Amyloid beta peptide-(1-42) induces internalization and degradation of beta2 adrenergic receptors in prefrontal cortical neurons. The Journal of biological chemistry 286, 31852–31863 (2011).

10. Udan, M.L., Ajit, D., Crouse, N.R. & Nichols, M.R. Toll-like receptors 2 and 4 mediate Abeta(1-42) activation of the innate immune response in a human monocytic cell line. J Neurochem 104, 524–533 (2008).

11. Chaney, M.O., et al. RAGE and amyloid beta interactions: atomic force microscopy and molecular modeling. Biochimica et biophysica acta 1741, 199–205 (2005).

12. Tran, L., Basdevant, N., Prevost, C. & Ha-Duong, T. Structure of ring-shaped Abeta(4)(2) oligomers determined by conformational selection. Sci Rep 6, 21429 (2016).

13. Xue, C., et al. Abeta42 fibril formation from predominantly oligomeric samples suggests a link between oligomer heterogeneity and fibril polymorphism. R Soc Open Sci 6, 190179 (2019).

14. Bharadwaj, P., et al. Role of the cell membrane interface in modulating production and uptake of Alzheimer’s beta amyloid protein. Biochim Biophys Acta Biomembr 1860, 1639–1651 (2018).

15. Mark, R.J., Pang, Z., Geddes, J.W., Uchida, K. & Mattson, M.P. Amyloid beta-peptide impairs glucose transport in hippocampal and cortical neurons: involvement of membrane lipid peroxidation. J Neurosci 17, 1046–1054 (1997).

16. Prapong, T., et al. Amyloid beta-peptide decreases neuronal glucose uptake despite causing increase in GLUT3 mRNA transcription and GLUT3 translocation to the plasma membrane. Exp Neurol 174, 253–258 (2002).

17. Malkov, A., et al. Abeta initiates brain hypometabolism, network dysfunction and behavioral abnormalities via NOX2-induced oxidative stress in mice. Commun Biol 4, 1054 (2021).

18. Casley, C.S., Canevari, L., Land, J.M., Clark, J.B. & Sharpe, M.A. Beta-amyloid inhibits integrated mitochondrial respiration and key enzyme activities. J Neurochem 80, 91–100 (2002).

19. Moreira, P.I., Santos, M.S., Moreno, A. & Oliveira, C. Amyloid beta-peptide promotes permeability transition pore in brain mitochondria. Biosci Rep 21, 789–800 (2001).

20. Mossmann, D., et al. Amyloid-beta peptide induces mitochondrial dysfunction by inhibition of preprotein maturation. Cell metabolism 20, 662–669 (2014).

21. Tillement, L., Lecanu, L. & Papadopoulos, V. Further evidence on mitochondrial targeting of beta-amyloid and specificity of beta-amyloid-induced mitotoxicity in neurons. Neurodegener Dis 8, 331–344 (2011).

22. Xie, L., et al. Alzheimer’s beta-amyloid peptides compete for insulin binding to the insulin receptor. J Neurosci 22, RC221 (2002).

23. Roher, A.E., et al. Amyloid beta peptides in human plasma and tissues and their significance for Alzheimer’s disease. Alzheimer’s & dementia : the journal of the Alzheimer’s Association 5, 18–29 (2009).

24. Lee, Y.H., et al. Amyloid precursor protein expression is upregulated in adipocytes in obesity. Obesity (Silver Spring) 16, 1493–1500 (2008).

25. An, Y.A., et al. Dysregulation of Amyloid Precursor Protein Impairs Adipose Tissue Mitochondrial Function and Promotes Obesity. Nat Metab 1, 1243–1257 (2019).

26. Shigemori, K., Nomura, S., Umeda, T., Takeda, S. & Tomiyama, T. Peripheral Abeta acts as a negative modulator of insulin secretion. Proceedings of the National Academy of Sciences of the United States of America 119, e2117723119 (2022).

27. Balakrishnan, K., et al. Plasma Abeta42 correlates positively with increased body fat in healthy individuals. Journal of Alzheimer’s disease : JAD 8, 269–282 (2005).

28. Fujiwara, T., Oda, K., Yokota, S., Takatsuki, A. & Ikehara, Y. Brefeldin A causes disassembly of the Golgi complex and accumulation of secretory proteins in the endoplasmic reticulum. The Journal of biological chemistry 263, 18545–18552 (1988).

29. Lopaschuk, G.D., Folmes, C.D. & Stanley, W.C. Cardiac energy metabolism in obesity. Circulation research 101, 335–347 (2007).

30. Rayner, J.J., et al. The relative contribution of metabolic and structural abnormalities to diastolic dysfunction in obesity. International journal of obesity 42, 441–447 (2018).

31. Borlaug, B.A., et al. Obesity and heart failure with preserved ejection fraction: new insights and pathophysiologic targets. Cardiovascular research (2022).

32. Obokata, M., Reddy, Y.N.V., Pislaru, S.V., Melenovsky, V. & Borlaug, B.A. Evidence Supporting the Existence of a Distinct Obese Phenotype of Heart Failure With Preserved Ejection Fraction. Circulation 136, 6–19 (2017).

33. Koenig, A.L., et al. Single-cell transcriptomics reveals cell-type specific diversification in human heart failure. Nature Cardiovascular Research 1, 263–280 (2022).

34. Yan, Y. & Wang, C. Abeta42 is more rigid than Abeta40 at the C terminus: implications for Abeta aggregation and toxicity. Journal of molecular biology 364, 853–862 (2006).

35. Hoglund, K., et al. Plasma levels of beta-amyloid(1-40), beta-amyloid(1-42), and total beta-amyloid remain unaffected in adult patients with hypercholesterolemia after treatment with statins. Arch Neurol 61, 333–337 (2004).

36. Czeczor, J.K., et al. APP deficiency results in resistance to obesity but impairs glucose tolerance upon high fat feeding. J Endocrinol 237, 311–322 (2018).

37. Meakin, P.J., et al. Reduction in BACE1 decreases body weight, protects against diet-induced obesity and enhances insulin sensitivity in mice. The Biochemical journal 441, 285–296 (2012).

38. Zago, W., et al. Neutralization of soluble, synaptotoxic amyloid beta species by antibodies is epitope specific. J Neurosci 32, 2696–2702 (2012).

39. Salloway, S., et al. Two phase 3 trials of bapineuzumab in mild-to-moderate Alzheimer’s disease. The New England journal of medicine 370, 322–333 (2014).

40. Mureddu, G.F., de Simone, G., Greco, R., Rosato, G.F. & Contaldo, F. Left ventricular filling pattern in uncomplicated obesity. The American journal of cardiology 77, 509–514 (1996).

41. Aljaroudi, W., et al. Impact of body mass index on diastolic function in patients with normal left ventricular ejection fraction. Nutr Diabetes 2, e39 (2012).

42. Ingul, C.B., Tjonna, A.E., Stolen, T.O., Stoylen, A. & Wisloff, U. Impaired cardiac function among obese adolescents: effect of aerobic interval training. Arch Pediatr Adolesc Med 164, 852–859 (2010).

43. Gaur, V., et al. Scriptaid enhances skeletal muscle insulin action and cardiac function in obese mice. Diabetes, obesity & metabolism 19, 936–943 (2017).

44. Demattos, R.B., et al. A plaque-specific antibody clears existing beta-amyloid plaques in Alzheimer’s disease mice. Neuron 76, 908–920 (2012).

45. McFarland, T.M., Alam, M., Goldstein, S., Pickard, S.D. & Stein, P.D. Echocardiographic diagnosis of left ventricular hypertrophy. Circulation 57, 1140–1144 (1978).

46. Ndumele, C.E., et al. Obesity and Subtypes of Incident Cardiovascular Disease. J Am Heart Assoc 5(2016).

47. Hampel, H., et al. The Amyloid-beta Pathway in Alzheimer’s Disease. Mol Psychiatry 26, 5481–5503 (2021).

48. Kaspi, A. & Ziemann, M. mitch: multi-contrast pathway enrichment for multi-omics and single-cell profiling data. BMC Genomics 21, 447 (2020).

49. Boudina, S., et al. Reduced mitochondrial oxidative capacity and increased mitochondrial uncoupling impair myocardial energetics in obesity. Circulation 112, 2686–2695 (2005).

50. Chatham, J.C. & Seymour, A.M. Cardiac carbohydrate metabolism in Zucker diabetic fatty rats. Cardiovascular research 55, 104–112 (2002).

51. Lewis, A.J., Neubauer, S., Tyler, D.J. & Rider, O.J. Pyruvate dehydrogenase as a therapeutic target for obesity cardiomyopathy. Expert Opin Ther Targets 20, 755–766 (2016).

52. Chen, J.X. & Yan, S.D. Amyloid-beta-induced mitochondrial dysfunction. Journal of Alzheimer’s disease : JAD 12, 177–184 (2007).

53. Lustbader, J.W., et al. ABAD directly links Abeta to mitochondrial toxicity in Alzheimer’s disease. Science 304, 448–452 (2004).

54. Farris, W., et al. Insulin-degrading enzyme regulates the levels of insulin, amyloid beta-protein, and the beta-amyloid precursor protein intracellular domain in vivo. Proceedings of the National Academy of Sciences of the United States of America 100, 4162–4167 (2003).

55. Cole, S.L. & Vassar, R. The Alzheimer’s disease beta-secretase enzyme, BACE1. Mol Neurodegener 2, 22 (2007).

56. Troncone, L., et al. Abeta Amyloid Pathology Affects the Hearts of Patients With Alzheimer’s Disease: Mind the Heart. J Am Coll Cardiol 68, 2395–2407 (2016).

57. Mehta, P.D., Capone, G., Jewell, A. & Freedland, R.L. Increased amyloid beta protein levels in children and adolescents with Down syndrome. J Neurol Sci 254, 22–27 (2007).

58. Al-Biltagi, M., Serag, A.R., Hefidah, M.M. & Mabrouk, M.M. Evaluation of cardiac functions with Doppler echocardiography in children with Down syndrome and anatomically normal heart. Cardiol Young 23, 174–180 (2013).

59. Yousefirad, N., Kaygisiz, Z. & Aydin, Y. The Effects of Beta Amyloid Peptide 1-42 on Isolated Rat Hearts and Ileum Smooth Muscle. Pharmacology 98, 261–266 (2016).

60. Dabkowski, E.R., et al. Mitochondrial dysfunction in the type 2 diabetic heart is associated with alterations in spatially distinct mitochondrial proteomes. American journal of physiology. Heart and circulatory physiology 299, H529–540 (2010).

61. Andres, A.M., et al. Discordant signaling and autophagy response to fasting in hearts of obese mice: Implications for ischemia tolerance. American journal of physiology. Heart and circulatory physiology 311, H219–228 (2016).

62. Jang, S., Chapa-Dubocq, X.R., Parodi-Rullan, R.M., Fossati, S. & Javadov, S. Beta-Amyloid Instigates Dysfunction of Mitochondria in Cardiac Cells. Cells 11(2022).

63. Nazere, K., et al. Amyloid Beta Is Internalized via Macropinocytosis, an HSPG-and Lipid Raft-Dependent and Rac1-Mediated Process. Frontiers in Molecular Neuroscience 15(2022).

64. Lai, A.Y. & McLaurin, J. Mechanisms of Amyloid-Beta Peptide Uptake by Neurons: The Role of Lipid Rafts and Lipid Raft-Associated Proteins. Internation Journal of Alzheimers Disease 2011, 548380 (2011).

65. Dunlay, S.M., Roger, V.L. & Redfield, M.M. Epidemiology of heart failure with preserved ejection fraction. Nat Rev Cardiol 14, 591–602 (2017).

66. Fung, J., Frost, D., Chakrabartty, A. & McLaurin, J. Interaction of human and mouse Abeta peptides. J Neurochem 91, 1398–1403 (2004).

67. Venardos, K., De Jong, K.A., Elkamie, M., Connor, T. & McGee, S.L. The PKD Inhibitor CID755673 Enhances Cardiac Function in Diabetic db/db Mice. PloS one 10, e0120934 (2015).

68. De Jong, K.A., et al. Loss of protein kinase D activity demonstrates redundancy in cardiac glucose metabolism and preserves cardiac function in obesity. Molecular metabolism 42, 101105 (2020).

69. Lindsey, M.L., Kassiri, Z., Virag, J.A.I., de Castro Bras, L.E. & Scherrer-Crosbie, M. Guidelines for measuring cardiac physiology in mice. American journal of physiology. Heart and circulatory physiology 314, H733–H752 (2018).

70. Esparza, T.J., et al. Soluble Amyloid-beta Aggregates from Human Alzheimer’s Disease Brains. Sci Rep 6, 38187 (2016).

71. Martin, S.D., Morrison, S., Konstantopoulos, N. & McGee, S.L. Mitochondrial dysfunction has divergent, cell type-dependent effects on insulin action. Molecular metabolism 3, 408–418 (2014).

72. Wu, W., et al. Activation of Hippo signaling pathway mediates mitochondria dysfunction and dilated cardiomyopathy in mice. Theranostics 11, 8993–9008 (2021).

73. Williams, D., et al. Abnormal mitochondrial L-arginine transport contributes to the pathogenesis of heart failure and rexoygenation injury. PloS one 9, e104643 (2014).

74. Martin, S.D. & McGee, S.L. A systematic flux analysis approach to identify metabolic vulnerabilities in human breast cancer cell lines. Cancer & metabolism 7, 12 (2019).

75. Gaur, V., et al. Disruption of the Class IIa HDAC Corepressor Complex Increases Energy Expenditure and Lipid Oxidation. Cell reports 16, 2802–2810 (2016).

