## Supplementary information for "Amyloid beta 42 alters cardiac metabolism and impairs cardiac function in obesity"

This document contains 5 data tables and 6 data figures.

### Extended Data Table 1

| Parameter | ScrA$\beta_{42}$ | A$\beta_{42}$ | *P* value |
|---|---|---|---|
| Peak aortic flow (cm/sec) | 61.9 ± 3.8 | 58.7 ± 1.1 | 0.481 |
| Ejection time (msec) | 52.5 ± 1.6 | 50.3 ± 2.0 | 0.411 |
| Heart rate (bpm) | 386 ± 24 | 397 ± 18 | 0.736 |
| Peak E wave (cm/sec) | 58.9 ± 4.9 | 60.6 ± 2.7 | 0.770 |
| Peak A wave (cm/sec) | 27.4 ± 2.4 | 32.0 ± 0.6 | 0.123 |
| IVSd (cm) | 0.052 ± 0.001 | 0.053 ± 0.001 | 0.346 |
| LVIDd (cm) | 0.346 ± 0.006 | 0.346 ± 0.016 | 0.974 |
| LVPWd (cm) | 0.059 ± 0.001 | 0.057 ± 0.001 | 0.107 |
| IVSs (cm) | 0.062 ± 0.001 | 0.059 ± 0.001 | 0.086 |
| LVIDs (cm) | 0.230 ± 0.003 | 0.251 ± 0.013 | 0.151 |
| LVPWs (cm) | 0.072 ± 0.001 | 0.067 ± 0.001 | <0.001 |
| Estimated LV mass (mg) | 100.4 ± 3.4 | 90.0 ± 3.4 | 0.046 |
| Heart weight/tibia length (mg/mm) | 7.11 ± 0.09 | 7.19 ± 0.17 | 0.751 |

**Extended Data Table 1: Cardiac function and morphology in mice administered ScrA$\beta_{42}$ or A$\beta_{42}$.** IVSd, intraventricular septum thickness at diastole; LVIDd, left ventricular internal diameter at diastole; LVPWd, left ventricular posterior wall thickness at diastole; IVSs, intraventricular septum thickness at systole; LVIDs, left ventricular internal diameter at systole; LVPWs, left ventricular posterior wall thickness at systole. Data are mean ± SEM, n =10 mice/group. Groups compared by unpaired t-test, two-tailed.

### Extended Data Table 2

| Parameter | ScrAβ$_{40}$ | Aβ$_{40}$ | *P* value |
|---|---|---|---|
| Peak aortic flow (cm/sec) | 60.4 ± 4.2 | 66.2 ± 5.1 | 0.391 |
| Ejection time (msec) | 59.3 ± 1.4 | 61.5 ± 1.2 | 0.233 |
| Heart rate (bpm) | 416 ± 12 | 384 ± 10 | 0.060 |
| Peak E wave (cm/sec) | 57.1 ± 3.6 | 58.1 ± 4.4 | 0.862 |
| Peak A wave (cm/sec) | 35.9 ± 1.8 | 32.4 ± 1.8 | 0.193 |
| E:A ratio | 1.54 ± 0.08 | 1.73 ± 0.10 | 0.153 |
| Deceleration time (msec) | 0.032 ± 0.001 | 0.032 ± 0.001 | 0.939 |
| IVSd (cm) | 0.125 ± 0.005 | 0.120 ± 0.006 | 0.498 |
| LVIDd (cm) | 0.340 ± 0.010 | 0.350 ± 0.012 | 0.528 |
| LVPWd (cm) | 0.125 ± 0.004 | 0.128 ± 0.003 | 0.533 |
| IVSs (cm) | 0.178 ± 0.007 | 0.174 ± 0.007 | 0.697 |
| LVIDs (cm) | 0.225 ± 0.012 | 0.226 ± 0.013 | 0.981 |
| LVPWs (cm) | 0.141 ± 0.004 | 0.152 ± 0.005 | 0.094 |
| Ejection fraction (%) | 71.5 ± 2.6 | 73.2 ± 2.1 | 0.602 |
| Fractional shortening (%) | 35.9 ± 2.2 | 36.2 ± 1.7 | 0.905 |
| Estimated LV mass (mg) | 217 ± 15 | 204 ± 11 | 0.502 |
| Heart weight/tibia length (mg/mm) | 9.60 ± 0.42 | 8.78 ± 0.24 | 0.112 |

**Extended Data Table 2: Cardiac function and morphology in mice administered ScrAβ$_{40}$ or Aβ$_{40}$.** IVSd, intraventricular septum thickness at diastole; LVIDd, left ventricular internal diameter at diastole; LVPWd, left ventricular posterior wall thickness at diastole; IVSs, intraventricular septum thickness at systole; LVIDs, left ventricular internal diameter at systole; LVPWs, left ventricular posterior wall thickness at systole. Data are mean ± SEM, n =12 mice/group. Groups compared by unpaired t-test, two-tailed.

### Extended Data Table 3

| | Control | | 3D6 | | 2-way RM ANOVA *P* value | | |
|---|---|---|---|---|---|---|---|
| **Parameter** | **Pre** | **Post** | **Pre** | **Post** | **Tx** | **Time** | **Int.** |
| Peak aortic flow (cm/sec) | 72.2 ± 6.9 | 67.9 ± 4.7 | 67.0 ± 6.0 | 65.3 ± 4.7 | 0.5022 | 0.5975 | 0.8217 |
| Ejection time (msec) | 48.5 ± 2.3 | 54.0 ± 2.2 | 49.4 ± 3.2 | 52.5 ± 2.4 | 0.9121 | 0.0969 | 0.6365 |
| Heart rate (bpm) | 455 ± 22 | 453 ± 19 | 437 ± 17 | 475 ± 22 | 0.9328 | 0.3786 | 0.3177 |
| Peak E wave (cm/sec) | 48.3 ± 2.7 | 51.0 ± 3.8 | 51.1 ± 4.6 | 49.4 ± 3.3 | 0.8070 | 0.7936 | 0.3434 |
| Peak A wave (cm/sec) | 28.7 ± 1.6 | 37.4 ± 3.5 | 32.7 ± 3.1 | 35.0 ± 2.8 | 0.9631 | 0.1365 | 0.1724 |
| E:A ratio | 1.69 ± 0.08 | 1.41 ± 0.04 | 1.60 ± 0.09 | 1.32 ± 0.04 | 0.1654 | 0.0056 | 0.8433 |
| IVSd (cm) | 0.116 ± 0.003 | 0.128 ± 0.006 | 0.130 ± 0.006 | 0.129 ± 0.006 | 0.2207 | 0.1432 | 0.0829 |
| LVIDd (cm) | 0.272 ± 0.012 | 0.307 ± 0.009 | 0.272 ± 0.010 | 0.293 ± 0.013 | 0.3505 | 0.0072 | 0.7772 |
| LVPWd (cm) | 0.120 ± 0.006 | 0.135 ± 0.008 | 0.130 ± 0.007 | 0.131 ± 0.005 | 0.9798 | 0.0013 | 0.2343 |
| IVSs (cm) | 0.165 ± 0.002 | 0.174 ± 0.006 | 0.168 ± 0.007 | 0.166 ± 0.006 | 0.6910 | 0.5835 | 0.3740 |
| LVIDs (cm) | 0.175 ± 0.012 | 0.189 ± 0.010 | 0.177 ± 0.011 | 0.195 ± 0.012 | 0.7513 | 0.1800 | 0.8458 |
| LVPWs (cm) | 0.137 ± 0.006 | 0.158 ± 0.009 | 0.146 ± 0.005 | 0.148 ± 0.010 | 0.9417 | 0.1486 | 0.2658 |
| Ejection fraction (%) | 75.3 ± 1.5 | 75.0 ± 1.8 | 76.1 ± 1.8 | 69.5 ± 2.8 | 0.1797 | 0.3923 | 0.5001 |
| Fractional shortening (%) | 37.7 ± 1.3 | 37.6 ± 1.6 | 37.3 ± 1.9 | 34.6 ± 2.4 | 0.7223 | 0.1921 | 0.1953 |

**Extended Data Table 3: Cardiac function and morphology in mice fed a high fat diet and administered control or 3D6 antibodies.** IVSd, intraventricular septum thickness at diastole; LVIDd, left ventricular internal diameter at diastole; LVPWd, left ventricular posterior wall thickness at diastole; IVSs, intraventricular septum thickness at systole; LVIDs, left ventricular internal diameter at systole; LVPWs, left ventricular posterior wall thickness at systole. Data are mean ± SEM, n =12 mice/group. Groups compared by two-way repeated measures ANOVA. Tx, treatment; Int, interaction.

### Extended Data Table 4

| | Chow control | | | HFD control | | | HFD 3D6 | | | Mixed effects model *P* value | | |
|---|---|---|---|---|---|---|---|---|---|---|---|---|
| | Base-line | Pre-treat | Post-treat | Base-line | Pre-treat | Post-treat | Base-line | Pre-treat | Post-treat | Tx | Time | Int. |
| PAF (cm/sec) | 57.8 ± 2.7 | 59.1 ± 4.2 | 61.3 ± 2.6 | 60.1 ± 6.3 | 59.3 ± 3.2 | 67.7 ± 5.1 | 58.7 ± 4.0 | 64.1 ± 3.0 | 71.8 ± 2.5 | 0.0955 | 0.0067 | 0.5443 |
| ET (msec) | 42.3 ± 1.9 | 42.3 ± 2.2 | 46.8 ± 2.0 | 41.2 ± 1.2 | 49.2 ± 2.4 | 43.8 ± 3.2 | 44.5 ± 1.5 | 46.2 ± 1.5 | 45.3 ± 2.6 | 0.6138 | 0.0318 | 0.5453 |
| HR (bpm) | 468 ± 18 | 491 ± 17 | 510 ± 23 | 441 ± 4 | 501 ± 9 | 556 ± 23 | 462 ± 10 | 503 ± 12 | 572 ± 12 | 0.3321 | < 0.0001 | 0.3321 |
| Peak E (cm/sec) | 42.3 ± 3.5 | 51.1 ± 3.8 | 50.0 ± 4.1 | 45.2 ± 4.2 | 58.3 ± 4.1 | 53.0 ± 2.6 | 41.6 ± 1.4 | 52.3 ± 2.8 | 63.8 ± 3.5 | 0.2964 | 0.0003 | 0.2945 |
| Peak A (cm/sec) | 27.3 ± 2.2 | 33.5 ± 2.7 | 35.0 ± 1.9 | 24.4 ± 1.2 | 34.4 ± 2.4 | 36.2 ± 2.8 | 28.7 ± 1.7 | 32.3 ± 2.2 | 46.7 ± 3.8 | 0.0852 | < 0.0001 | 0.0592 |
| E:A ratio | 1.62 ± 0.08 | 1.59 ± 0.03 | 1.57 ± 0.03 | 1.65 ± 0.05 | 1.67 ± 0.06 | 1.45 ± 0.09 | 1.52 ± 0.05 | 1.56 ± 0.07 | 1.41 ± 0.06 | 0.2417 | 0.1167 | 0.8675 |
| IVSd (cm) | 0.147 ± 0.002 | 0.109 ± 0.005 | 0.118 ± 0.004 | 0.135 ± 0.006 | 0.114 ± 0.003 | 0.129 ± 0.005 | 0.137 ± 0.008 | 0.125 ± 0.005 | 0.122 ± 0.003 | 0.7234 | < 0.0001 | 0.0335 |
| LVIDd (cm) | 0.248 ± 0.011 | 0.314 ± 0.014 | 0.293 ± 0.014 | 0.268 ± 0.005 | 0.311 ± 0.015 | 0.308 ± 0.015 | 0.263 ± 0.014 | 0.303 ± 0.014 | 0.281 ± 0.009 | 0.4968 | 0.0001 | 0.6252 |
| LVPWd (cm) | 0.156 ± 0.008 | 0.136 ± 0.008 | 0.125 ± 0.008 | 0.142 ± 0.005 | 0.152 ± 0.007 | 0.137 ± 0.009 | 0.134 ± 0.007 | 0.147 ± 0.008 | 0.144 ± 0.007 | 0.6857 | 0.0577 | 0.0241 |
| IVSs (cm) | 0.189 ± 0.008 | 0.158 ± 0.006 | 0.162 ± 0.006 | 0.175 ± 0.008 | 0.169 ± 0.005 | 0.178 ± 0.006 | 0.167 ± 0.009 | 0.169 ± 0.003 | 0.173 ± 0.004 | 0.9053 | 0.0876 | 0.0313 |
| LVIDs (cm) | 0.160 ± 0.007 | 0.193 ± 0.013 | 0.185 ± 0.015 | 0.192 ± 0.010 | 0.192 ± 0.016 | 0.185 ± 0.015 | 0.186 ± 0.012 | 0.188 ± 0.014 | 0.172 ± 0.010 | 0.4420 | 0.6350 | 0.6135 |
| LVPWs (cm) | 0.171 ± 0.005 | 0.156 ± 0.007 | 0.141 ± 0.008 | 0.160 ± 0.006 | 0.174 ± 0.010 | 0.166 ± 0.011 | 0.149 ± 0.009 | 0.167 ± 0.008 | 0.157 ± 0.008 | 0.2567 | 0.2109 | 0.0370 |
| EF (%) | 71.0 ± 1.9 | 76.1 ± 2.1 | 74.1 ± 2.9 | 67.3 ± 2.6 | 74.9 ± 3.0 | 78.2 ± 2.2 | 66.2 ± 1.6 | 74.3 ± 3.1 | 76.7 ± 2.2 | 0.8585 | 0.004 | 0.5063 |
| FS (%) | 34.0 ± 1.4 | 39.0 ± 2.1 | 37.5 ± 2.6 | 31.8 ± 1.8 | 39.5 ± 3.0 | 40.9 ± 2.3 | 30.0 ± 0.9 | 38.3 ± 2.9 | 39.4 ± 2.0 | 0.9689 | 0.0005 | 0.6390 |
| LV mass (mg) | 152 ± 7 | 153 ± 9 | 136 ± 3 | 161 ± 10 | 164 ± 5 | 161 ± 8 | 145 ± 14 | 166 ± 6 | 153 ± 5 | 0.0925 | 0.1080 | 0.1357 |

**Extended Data Table 4: Cardiac function and morphology in mice fed chow or high fat diet and administered control or 3D6 antibodies.** PAF, peak aortic flow; ET, ejection time; HR, heart rate; IVSd, intraventricular septum thickness at diastole; LVIDd, left ventricular internal diameter at diastole; LVPWd, left ventricular posterior wall thickness at diastole; IVSs, intraventricular septum thickness at systole; LVIDs, left ventricular internal diameter at systole; LVPWs, left ventricular posterior wall thickness at systole. EF, ejection fraction; FS, fractional shortening. Data are mean ± SEM, n =12 mice/group. Groups compared by mixed effects model. Tx, treatment; Int, interaction.

### Extended Data Table 5

| Pathway | setSize | p MANOVA | p.adjust MANOVA | s.dist | s.A$\beta_{42}$ | s.3D6 | p.A$\beta_{42}$ | p.3D6 |
|---|---|---|---|---|---|---|---|---|
| TCA cycle | 22 | <0.0001 | 0.0004 | 0.608 | -0.350 | 0.498 | 0.0045 | <0.0001 |
| Pyruvate metabolism and TCA cycle | 50 | <0.0001 | <0.0001 | 0.567 | -0.305 | 0.478 | 0.0002 | <0.0001 |
| Pyruvate metabolism | 26 | 0.0002 | 0.0030 | 0.484 | -0.246 | 0.417 | 0.0301 | 0.0002 |
| Mitochondrial biogenesis | 71 | 0.0009 | 0.0014 | 0.268 | -0.184 | 0.196 | 0.0076 | 0.0044 |
| Protein localisation | 141 | 0.0001 | 0.0004 | 0.215 | -0.146 | 0.158 | 0.0028 | 0.0012 |
| Neddylation | 201 | <0.0001 | <0.0001 | 0.199 | -0.109 | 0.166 | 0.0079 | <0.0001 |
| Antigen processing | 262 | <0.0001 | <0.0001 | 0.197 | -0.075 | 0.182 | 0.0380 | <0.0001 |
| Autophagy | 117 | 0.0037 | 0.0050 | 0.188 | -0.127 | 0.139 | 0.0178 | 0.0097 |
| Chromatin modifying enzymes | 188 | 0.0006 | 0.0011 | 0.171 | -0.111 | 0.130 | 0.0002 | 0.0022 |
| Chromatin organisation | 188 | 0.0006 | 0.0011 | 0.171 | -0.111 | 0.130 | 0.0002 | 0.0022 |
| Hemostasis | 373 | <0.0001 | <0.0001 | 0.184 | 0.094 | -0.157 | 0.0019 | <0.0001 |
| Platelet activation | 187 | <0.0001 | <0.0001 | 0.214 | 0.098 | -0.190 | 0.0208 | <0.0001 |
| Elevated platelet cytosolic-Ca2+ | 97 | 0.0002 | 0.0005 | 0.252 | 0.148 | -0.203 | 0.0117 | 0.0005 |
| Platelet degranulation | 93 | 0.0001 | 0.0003 | 0.269 | 0.158 | -0.218 | 0.0084 | 0.0003 |
| Chondroitin sulfate metabolism | 40 | 0.0040 | 0.0050 | 0.319 | 0.205 | -0.244 | 0.0076 | 0.0044 |
| Kainate receptor activation | 20 | 0.0052 | 0.0060 | 0.439 | 0.339 | -0.278 | 0.0086 | 0.0314 |
| Basigin interactions | 18 | 0.0042 | 0.0291 | 0.473 | 0.343 | -0.326 | 0.0118 | 0.0168 |
| Thrombin signalling through PARs | 21 | 0.0013 | 0.0116 | 0.480 | 0.336 | -0.343 | 0.0076 | 0.0065 |
| Gβχ signalling through BTK | 11 | 0.0029 | 0.0021 | 0.508 | 0.371 | -0.346 | 0.00331 | 0.0468 |

**Extended Data Table 5: Reactome pathways reciprocally regulated in mice administered A$\beta_{42}$ or 3D6 relative to their respective control groups, as determined by mitch.**

**Extended Data Figure 1**

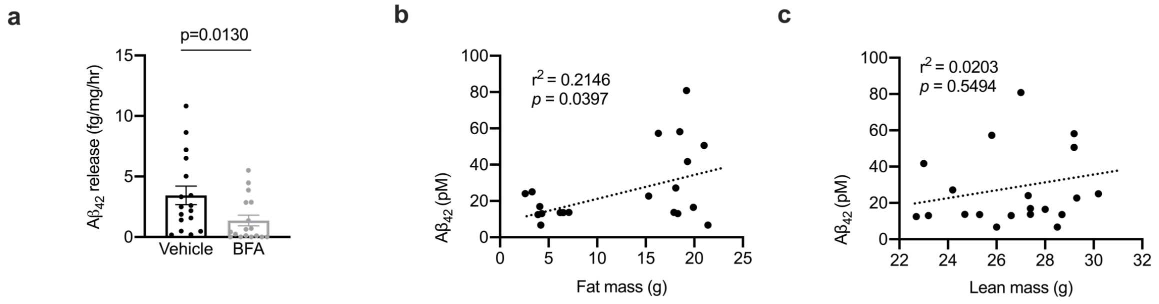


**Extended Data Figure 1: Aβ$_{42}$ release by adipose tissue and body composition correlations with plasma Aβ$_{42}$. a,** relative release of Aβ$_{42}$ from adipose tissue exposed to vehicle or Brefeldin A (BFA; Mann-Whitney test, U=73). **b**, correlation between fat mass and plasma Aβ$_{42}$. **c**, Correlation between lean mass and plasma Aβ$_{42}$. Two-tailed Pearsons correlation coefficient test. Data are mean ± SEM, n = 16 explants/group, 18 mice/correlation. Statistical tests are two-tailed.

**Extended Data Figure 2**

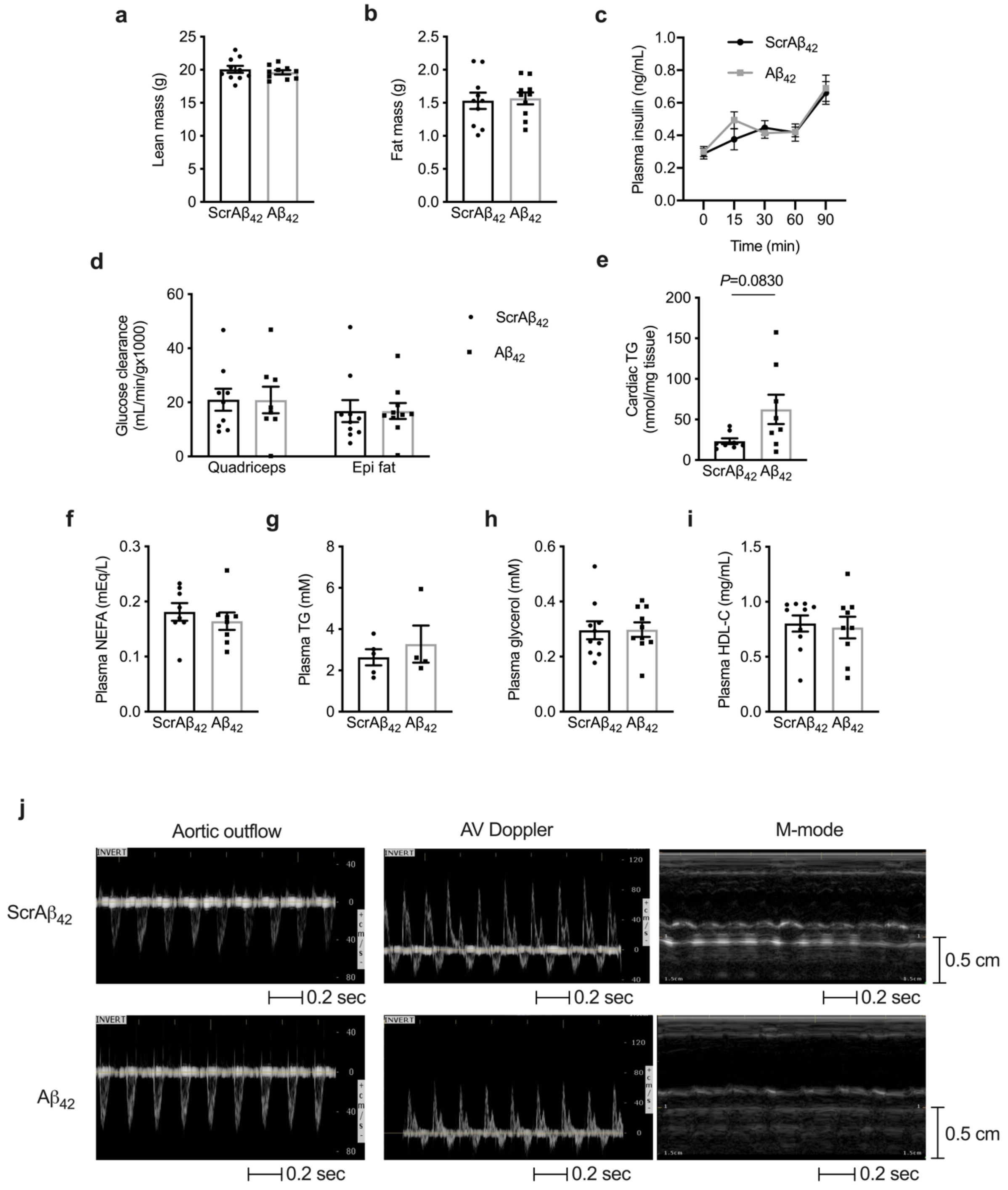


**Extended Data Figure 2: Characterisation of mice administered ScrAβ42 or Aβ42. a**, lean mass; **b**, fat mass; **c**, plasma insulin during a glucose tolerance test; **d**, glucose clearance by the quadriceps skeletal muscle and epididymal fat pad; **e**, $^{14}$C-glucose incorporation into cardiac lipids; **f**, plasma non-esterified fatty acids (NEFA); **g**, plasma triglycerides (TG); **h**, plasma glycerol; **i**, plasma high-density lipoprotein cholesterol (HDL-C), and; **j**, representative echocardiography images in mice administered ScrAβ42 or Aβ42. Data are mean ± SEM, n =6-10 mice/group.

### Extended Data Figure 3

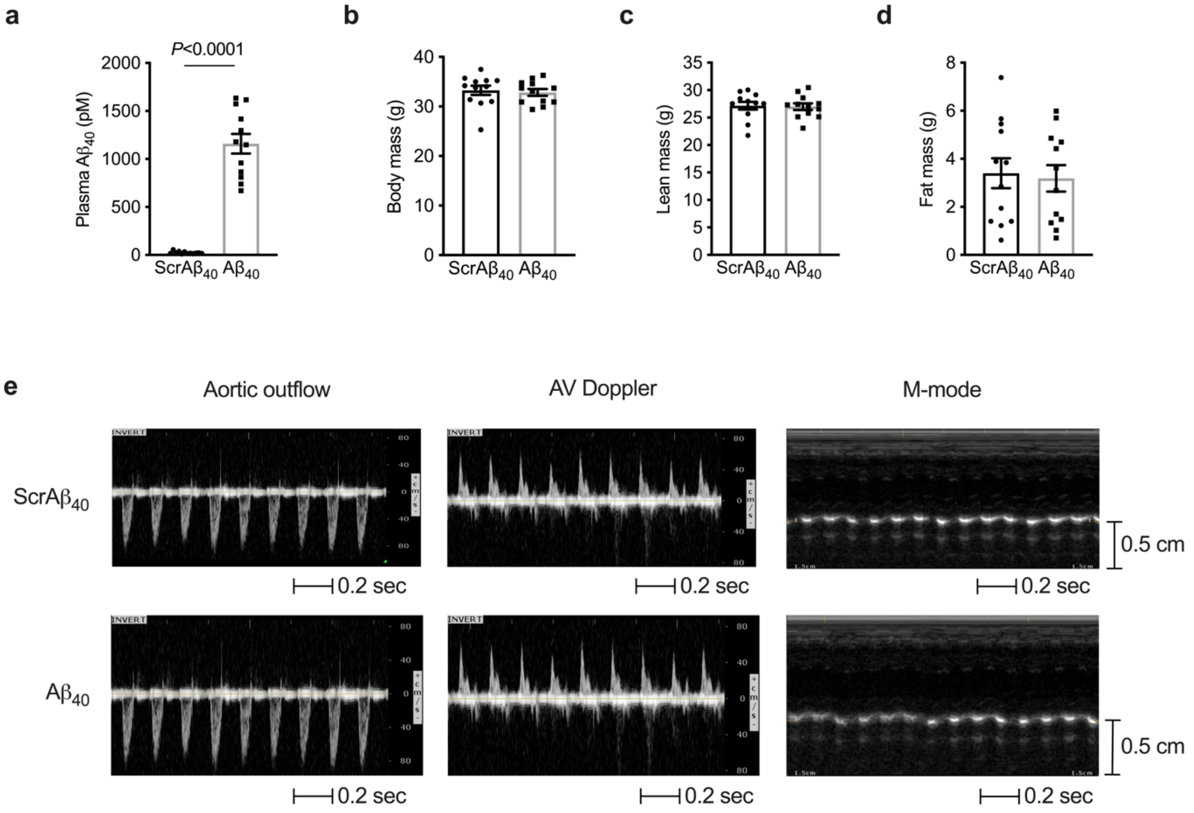


**Extended Data Figure 3: Characterisation of mice administered ScrA$\beta_{40}$ or A$\beta_{40}$. a**, plasma A$\beta_{40}$ 60 min after A$\beta_{40}$ administration. **b**, body weight; **c**, lean mass; **d**, fat mass, and; **e**, representative echocardiography images in mice administered ScrA$\beta_{40}$ or A$\beta_{40}$. Data are mean ± SEM, n =12 mice/group.

### Extended Data Figure 4

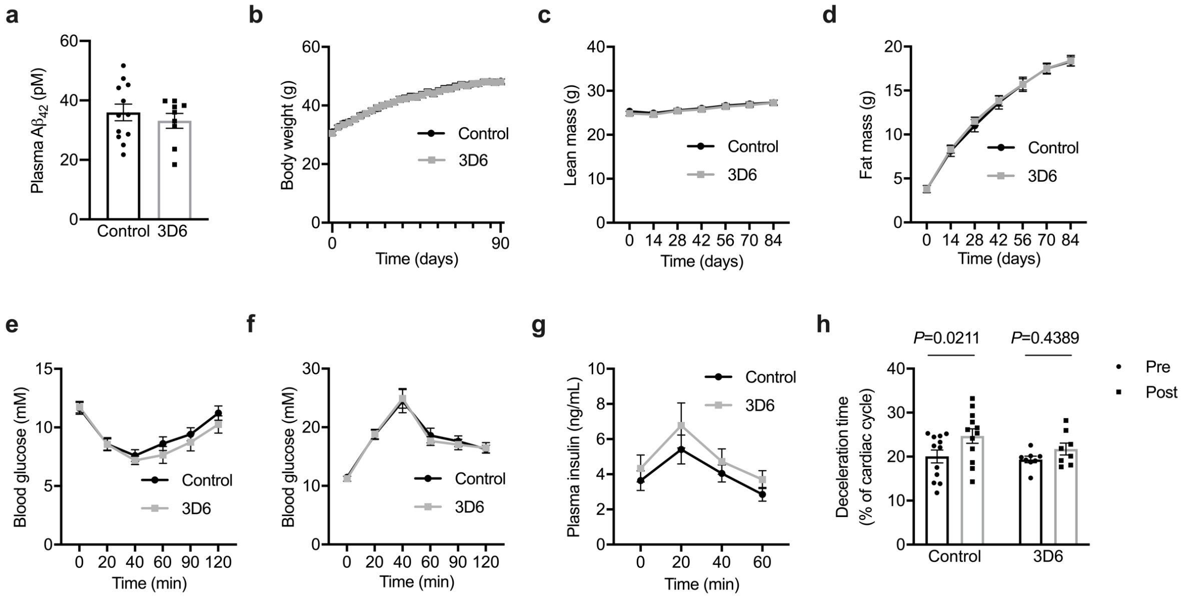


**Extended Data Figure 4: Characterisation of mice fed a high fat diet and administered control or 3D6 antibodies. a**, plasma A$\beta_{42}$; **b**, body weight; **c**, lean mass; **d**, fat mass, **e**, blood glucose during the insulin tolerance test; **f**, blood glucose during the glucose tolerance test; **g**, plasma insulin during the glucose tolerance test, and; **h**, cardiac deceleration time expressed as a percentage of the cardiac cycle (mixed effects model (time $P = 0.0140$, $F(1,18) = 7.407$) with Sidak's repeated measures test *P*.adjusted). Data are mean ± SEM, n =12 mice/group.

### Extended Data Figure 5

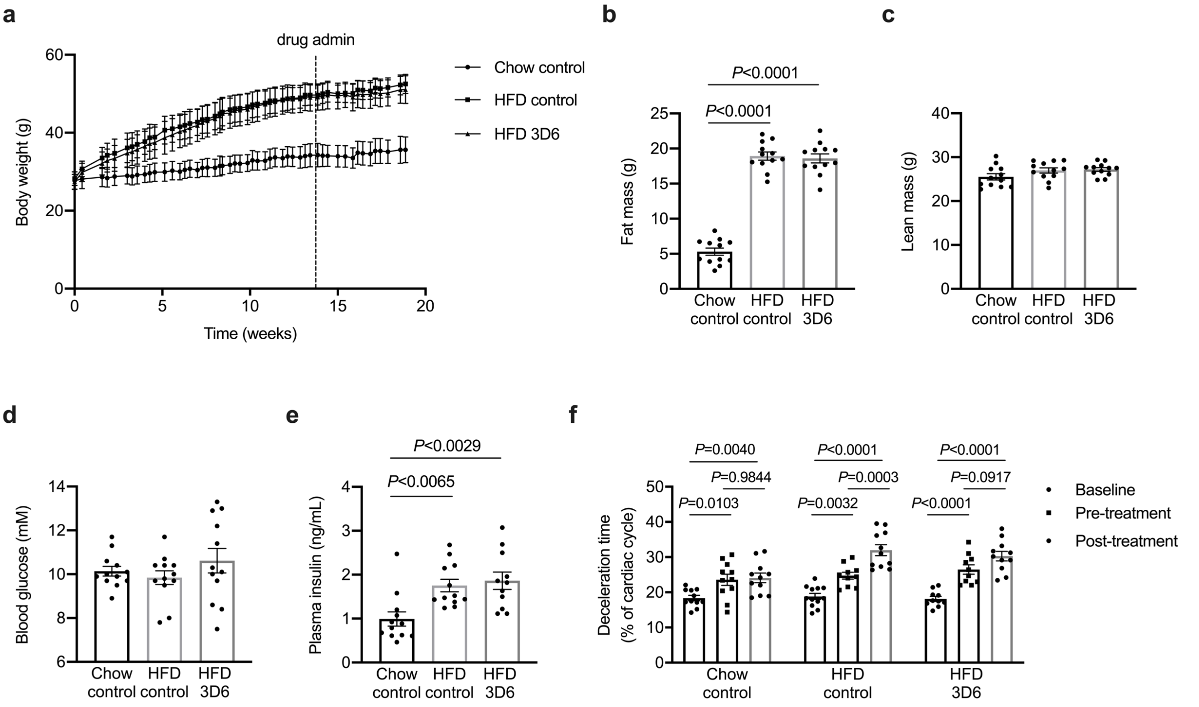


**Extended Data Figure 5: Characterisation of mice fed a high fat diet and administered control or 3D6 antibodies.** **a**, body weight; **b**, fat mass (one-way ANOVA ($P < 0.0001$; F(2,33) = 177.6) with Sidak's repeated measures test *P*.adjusted); **c**, lean mass, **d**, fasting blood glucose; **e**, fasting plasma insulin (Kruskal-Wallis test ($P = 0.0011$; $X^2 = 13.66$) with Dunn's repeated measures test *P*.adjusted), and; **h**, cardiac deceleration time expressed as a percentage of the cardiac cycle (mixed effects model (time $P < 0.0001$, $F(2,56) = 56.37$; treatment $P = 0.0044$, $F(2,32) = 6.445$; interaction $P = 0.0145$, $F(4,56) = 3.410$) with Sidak's repeated measures test *P*.adjusted). Data are mean ± SEM, n =12 mice/group.

### Extended Data Figure 6

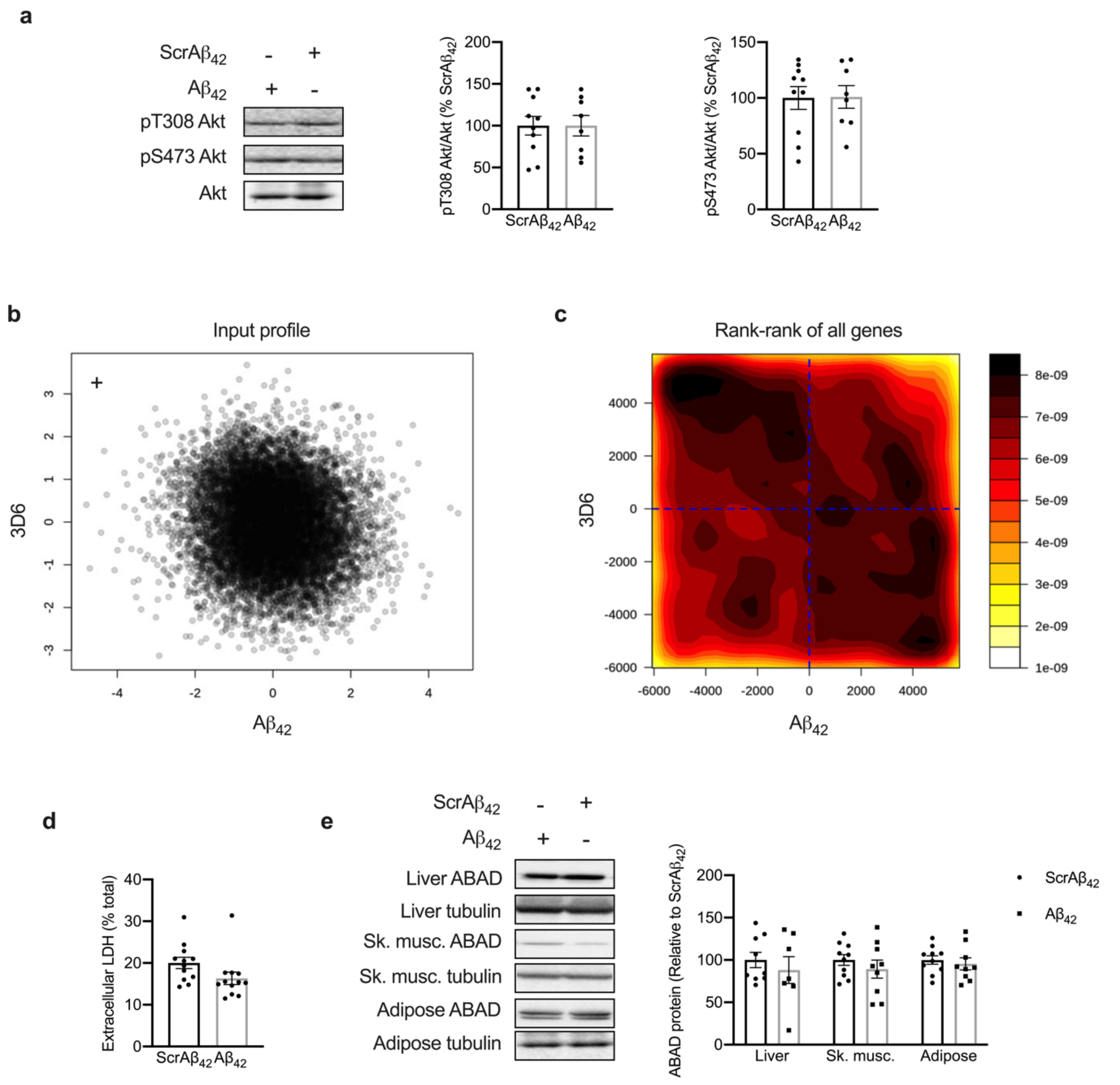


**Extended Data Figure 6: a**, phosphorylated T308 and S473 Akt in hearts of mice administered ScrA$\beta_{42}$ or A$\beta_{42}$ immediately following a glucose tolerance test. **b**, input profile defined as differential expression score defined as the sign of the fold change multiplied by the -log10(p-value), and; **c**, gene ranks from bulk RNA-seq analysis of gene expression in the hearts of mice administered A$\beta_{42}$ compared with mice administered ScrA$\beta_{42}$ (x-axis) and hearts of mice administered 3D6 antibody compared with mice administered control antibody (y-axis). **d,** extracellular lactate dehydrogenase (LDH) in primary neonatal ventricular cardiomyocytes (NVCM) exposed to ScrAb$_{42}$ or Ab$_{42}$ for 48 hr. **e,** amyloid binding alcohol dehydrogenase (ABAD) abundance in liver, skeletal muscle (Sk. musc.) and adipose tissue of mice administered ScrA$\beta_{42}$ or A$\beta_{42}$. Data are mean ± SEM, n=7-9 mice/group for protein analyses, n=6 mice/group for RNA seq analyses and 12-14 biological replicates/group for *in vitro* studies.
